# Human milk-derived extracellular vesicles promote the heat shock response in polarized microglia

**DOI:** 10.1101/2025.01.06.631449

**Authors:** Jasmyne A. Storm, Jueqin Lu, Mon Francis Obtial, Sanoji Wijenayake

## Abstract

Proteotoxic stress induces microglia polarization and attenuates cytoprotective, pro-survival cellular cascades. Milk-derived extracellular vesicles (MEVs) are lipid-coated nanovesicles that combat pro-inflammation in peripheral cells and tissues; however, the cytoprotective potential of MEVs remains unknown in brain macrophages. We investigated whether MEVs reduce neuroinflammation in human microglia by activating the heat shock response (HSR). The HSR triggers the upregulation of molecular chaperones (heat shock proteins; HSPs) to restore proteostasis by refolding or degrading misfolded aggregates. MEVs were isolated from unpasteurized human donor milk. Human microglia clone 3 (HMC3) cells were primed with 10 ng/mL IFN-γ to induce polarization, and a subset of cells were supplemented with 200 µg of MEVs. The abundance of HSF1 and candidate HSPs (Hsp70, Hsp90, Hsp40, Hsp27) were analyzed via RT-qPCR and western immunoblotting at 6h, 12h, and 24h post-MEV supplementation. We found that MEV supplementation promoted the HSR in polarized microglia, compared to homeostatic cells. Furthermore, MEVs increased the duration of the HSR in response to pro-inflammatory stress, exerting robust and continued pro-survival benefits.

## 1.0 Introduction

The Developmental Origins of Health and Disease (DOHaD) hypothesis describes how exposure to proteotoxic stress during perinatal life, the collective prenatal and postnatal periods, can influence development and disease outcomes in offspring [1–4]. Examples of potent perinatal stress that can mediate offspring programming include maternal diet, immune activation, maternal psychosocial state, substance abuse, and environmental exposures [2, 4–6]. One of the principal outcomes of perinatal stress is chronic systemic pro-inflammation and neuroinflammation [7]. Specifically, neuroinflammation that stems from microglia polarization and the attenuation of cytoprotective, pro-survival cellular cascades [7–9].

Microglia are the primary immune cells of the brain parenchyma and contribute to innate and adaptive immune responses and maintain a healthy central nervous system (CNS) [7, 10]. Microglia are involved in neurodevelopment and neurogenesis, synaptic pruning, phagocytosis, autophagy, and the secretion of cytokines, growth factors, and neurotrophic factors into the surrounding environment [11, 12]. Microglia are highly sensitive to stress and stimuli, which causes an enlargement of the soma, a shortening of the processes, and conversion into ameboid and rod shape cells [10, 13, 14]. While previously described as having distinct classic “M1” or alternate “M2” states, microglia polarization is more dynamic [15, 16]. Polarized microglia may produce an exaggerated response to inflammatory stimuli, exerting pro-inflammatory and/or anti-inflammatory effects in the CNS, which often result in changes in antigen presentation, cell proliferation, and/or morphology [17, 18].

Recent studies indicate that human milk promotes offspring development and provides pro-survival benefits through the transmission of bioactive, regulatory molecules during critical periods of early life. Specifically, human milk feeding could potentially attenuate the negative outcomes of select gestational stress [5, 19–25]. Milk is the primary nutritional source for most newborn mammals and is composed of carbohydrates, lipids, proteins, oligosaccharides, minerals, and vitamins [20, 26]. In addition to its nutritional components, milk contains bioactive molecules that shape offspring development, including hormones, growth factors, antibodies, stem cells, enzymes, peptides, bacteria that establish the gut microbiome, and milk-derived extracellular vesicles (MEVs) [5, 20, 27–29]. MEVs are lipid-coated nanovesicles, ranging from 30 to 150 nm in diameter, that are secreted primarily by mammary gland epithelial cells [30, 31]. MEVs have a lipid bilayer which protects the encapsulated cargo against gastrointestinal degradation and low pH environments. MEVs also travel via endo and transcytosis across complex biological barriers, including the intestinal endothelium and the blood brain barrier [19, 20, 32–35].

Recent studies have shown MEVs to be immunomodulatory and anti-inflammatory [36]. The primary function of MEVs is to deliver their cargo, including DNA, lipids, proteins, peptides, and small, microRNA (miRNA) to cells and tissues within the offspring’s system [30, 33, 37]. MEV miRNA are the most likely candidates to induce cellular regulation in offspring [38–40]. miRNA induce post-transcriptional regulation by binding to the 3’-UTR (untranslated region) of target mRNA and decrease or pause translation [41, 42]. This regulatory role allows MEV miRNA to modulate gene expression, cell-signalling, cell proliferation and differentiation, apoptosis, and immune regulation [34, 43–46]. MEVs have been shown to modulate numerous developmental pathways including epigenetic cascades, neurodevelopment, neurogenesis, and cell proliferation [19, 37, 45, 47, 48]. MEVs have also been shown to decrease rates of apoptosis *in vitro* and *in vivo* [44, 46]. Interestingly, MEVs exhibit cytoprotective, anti-inflammatory effects in peripheral tissues, including the intestinal epithelium, hepatocytes, and the lungs, where they attenuate nuclear factor kappa B (NFκB) mediated inflammation, and prevent the progression of inflammatory diseases, such as necrotizing enterocolitis [49–54]. The cytoprotective potential of MEVs in brain macrophages in response to neuroinflammation remains unknown. Specifically, the interaction between MEVs and existing pro-survival responses needs further investigation.

The heat shock response (HSR) is a critical baseline pro-survival response that protects cells from environmental or pathophysiological challenges (*e.g.*, exposure to cytokines/chemicals/toxins, heat, oxidative stress) [55, 56]. The HSR is regulated by DNA-binding heat shock-inducible transcription factors (HSFs) which regulate the transcription of molecular chaperones, called heat shock proteins (HSPs). HSPs prevent the aggregation of misfolded proteins and promote protein refolding or direct misfolded proteins to the proteosome for proteolytic degradation [57]. There are currently six known HSF1 isoforms, although not all have known DNA-binding domains [56]. Of these, HSF1 is the master regulator of the HSR in human and murine tissues, in addition to regulating the transcription of a broader variety of genes involved in the cell cycle, glucose metabolism, immune responses, autophagy, development and aging [56, 58–60].

In homeostatic cells, HSF1 is constitutively found in the cytoplasm in its inactive monomeric state, where it associates with Hsp70 and Hsp90 [61, 62]. During proteotoxic stress, HSF1 dissociates from Hsp70 and Hsp90 and trimerizes via the alignment of leucine zipper domains to allow a parallel orientation of each DNA-binding domain at the N-terminus [63, 64]. Following trimerization, HSF1 becomes hyperphosphorylated, translocates to the nucleus, and binds to the 5’-nGAAn-3’ sequence (also referred to as the AGAAN heat shock element) in the promotor region of target genes [60, 61, 65, 66]. HSF1 promotes the transcription of chaperones belonging to several HSP gene families, including candidate chaperones involved in protein refolding, including *HSPA* (*i.e.*, Hsp70), *HSPC* (*i.e.*, Hsp90), *DNAJ* (*i.e.*, Hsp40), and *HSPB* (small HSPs, *e.g.*, Hsp27) [55, 57].

Hsp70 and Hsp90 are both highly conserved ATP-dependent molecular chaperones, and are present in the cytoplasm, endoplasmic reticulum, and the mitochondria in eukaryotic cells [67]. Hsp70 and Hsp90 are binding partners, connected by the bridging co-chaperone Hop (Hsp-organizing protein) which together form the Hsp70-Hop-Hsp90 main refolding complex [62, 68–70]. In addition to their role in restoring proteostasis, the Hsp70-Hsp90 complex negatively regulates prolonged HSF1 activity [71]. A study by Kijima *et al*. (2018) [72] investigating Hsp90-HSF1 interactions in HEK293 cells, found that inhibition of the ATP-competitive, N-domain of Hsp90 disrupts its binding to HSF1, and prolongs HSF1 transcriptional activity. Furthermore, inhibition of Hsp70 and Hsp90 has been shown to increase HSF1 activation, suggesting that HSF1 activation is repressed in a feedback inhibition loop by Hsp70 and Hsp90 [72]. Hsp40 acts as a co-chaperone for Hsp70, binding misfolded protein clients and transferring them to Hsp70, as well as stimulating Hsp70 ATPase activity [73–75]. Hsp27 is an ATP-independent chaperone located in the cytoplasm, endoplasmic reticulum, and nucleus [76, 77]. Hsp27 is involved in the rapid disaggregation of misfolded proteins, preventing them from forming aggregates until ATP-dependent chaperones are available [59, 78–80].

The objective of this study is to further our understanding of the cytoprotective potential of MEVs, specifically in brain macrophages in response to an acute immune stress. We aim to investigate how MEVs may attenuate pro-inflammation and increase pro-survival responses, by exploring the relationship between MEVs and the HSR in human microglia. To our knowledge, this is the first study to explore the relationship between MEVs and baseline pro-survival pathways in the context of alleviating neuroinflammation and promoting neuroprotection.

## 2.0 Results

### 2.1 MEV characterization

We used nanoparticle tracking analysis (NTA) to determine MEV particle size and concentration. The isolated MEVs had a concentration of 2.13 x 10^11^ particles/mL with a mean size of 160 ± 0.5 nm (**Figure 1a**). MEV morphology was confirmed using transmission electron microscopy (TEM), allowing for distinct visualization of the electron-dense cargo, and the lipid bilayer outer membrane (**Figure 1b**). Some protein aggregates were detected alongside MEVs. No MEVs were detected in the negative control, although debris was detected. NTA and TEM findings confirm that the MEVs are within the typical 30 to 150 nm range. Western immunoblotting was used to detect the presence of two tetraspanin proteins (CD9, CD81) and a protein involved in EV biogenesis (syntenin-1), which were found in high abundance in the MEV lysates, with reduced levels found in the human microglia clone 3 (HMC3) cells (**Figure 1c**). Conversely, the negative control calnexin was absent from the MEV lysate but found in high abundance in the HMC3 cells. Calnexin, an endoplasmic reticulum marker that was used as the negative control to represent cellular protein distribution, was absent from the MEV lysate and abundant in HMC3 cells. The morphology and presence of positive endosome markers CD9, CD81, and syntenin-1, and distinct lack of the ER-marker calnexin, serve as a biological confirmation of our isolation and the lack of cellular contamination. Characterization using NTA, TEM, and western immunoblotting confirmed and validated our MEV isolates, as the observed morphology, size and detected biomarkers match the accepted standards published in accordance with MISEV [81].

**Figure 1.**
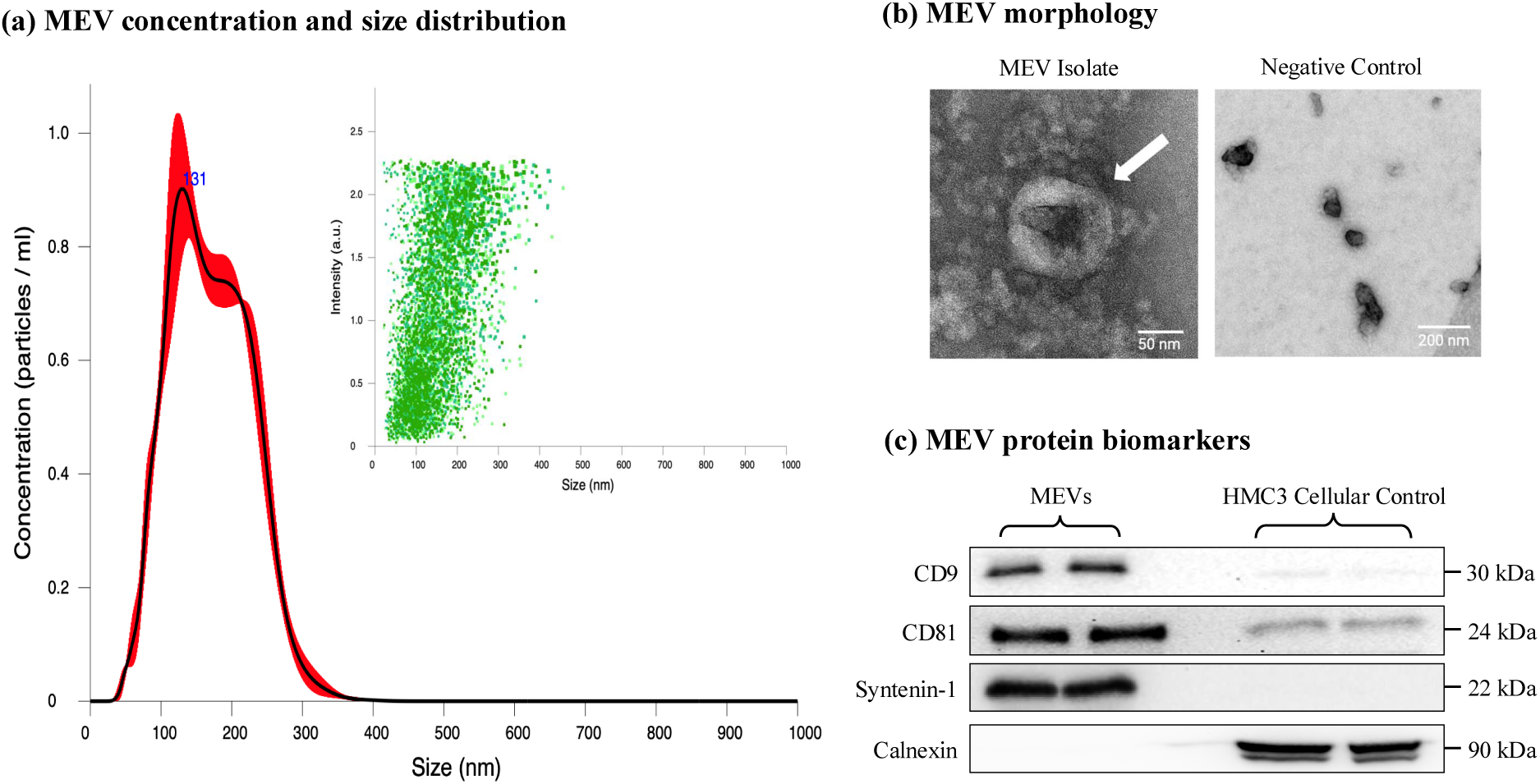
MEV characterization. **(a)** MEV concentration and size distribution were determined by nanoparticle tracking analysis (Malvern Instruments Ltd.; NanoSight NS300). MEVs were diluted 1:300 (v/v in PBS) and readings were recording using an sCMOS camera (level 15, detection threshold 10) using a 532nm green laser with 3x replicates at 30s capture speeds. **(b)** MEV morphology was determined by transmission electron microscopy (FEI Talos F200x S/TEM). MEVs were diluted 1:2 (v/v in PBS), loaded onto 400 mesh carbon-coated formvar film copper grids, and negatively stained using 2% uranyl acetate. **(c)** MEV protein biomarkers were detected using western immunoblotting. The endosome-specific transmembrane proteins CD9 and CD81, and the endosome-specific cytosolic protein syntenin-1 were used as positive controls. Calnexin, a marker of the endoplasmic reticulum, was used as a negative control to identify cellular protein contamination in the MEV isolates. Human microglia clone 3 (HMC3) was loaded as a cellular control.

### 2.2 Cell viability in response to priming and supplementation

To determine if priming via the pro-inflammatory cytokine interferon-gamma (IFN-γ) and/or MEV supplementation affected the viability of HMC3 cells, we conducted an MTT (3-[4,5- dimethylthiazol-2-yl]-2,5 diphenyl tetrazolium bromide) assay at 6, 12, and 24h (**Figure 2**). At 6h, cell viability remained unchanged across treatment groups (main effect of priming: (*F*(_1_,_16_) = 0.126, p = 0.727); main effect of supplementation: (*F*(_3_,_16_) = 0.057, p = 0.981)). Specifically, the cell viability (mean ± SEM) at 6h was 100.00% ± 3.50% for CTRL, 100.73% ± 7.58% for MEV, 97.21% ± 6.08% for PRI, and 98.86% ± 7.76% for PRI+MEV. At 12h, cell viability also remained unchanged across treatment groups (main effect of priming: (*F*(_1_,_15_) = 0.241, p = 0.630); main effect of supplementation: (*F*(_3_,_15_) = 1.027, p = 0.409)). The cell viability at 12h was 100.00% ± 6.68% for CTRL, 110.16% ± 3.89% for MEV, 105.53% ± 4.48% for PRI, and 95.48% ± 5.27% for PRI+MEV. At 24h, there was a main effect of priming (*F*(_1_,_14_) = 22.032, p < 0.001) where PRI cells (Tukey HSD, p = 0.015) and PRI+MEV (Tukey HSD, p = 0.005) cells had a lower cell viability compared to CTRL cells, and where PRI+MEV cells had a lower cell viability compared to MEV cells (Tukey HSD, p = 0.034). There is a main effect of time (*F*(_2,46_) = 11.644, p < 0.001), where cells at 24h had a lower cell viability compared to 6h cells (Tukey HSD, p = 0.002) and 12h cells (Tukey HSD, p < 0.001). The cell viability at 24h was 100.00% ± 4.85% for CTRL, 93.46% ± 2.81% for MEV, 74.20% ± 6.54% for PRI, and 68.12% ± 5.18% for PRI+MEV.

**Figure 2.**
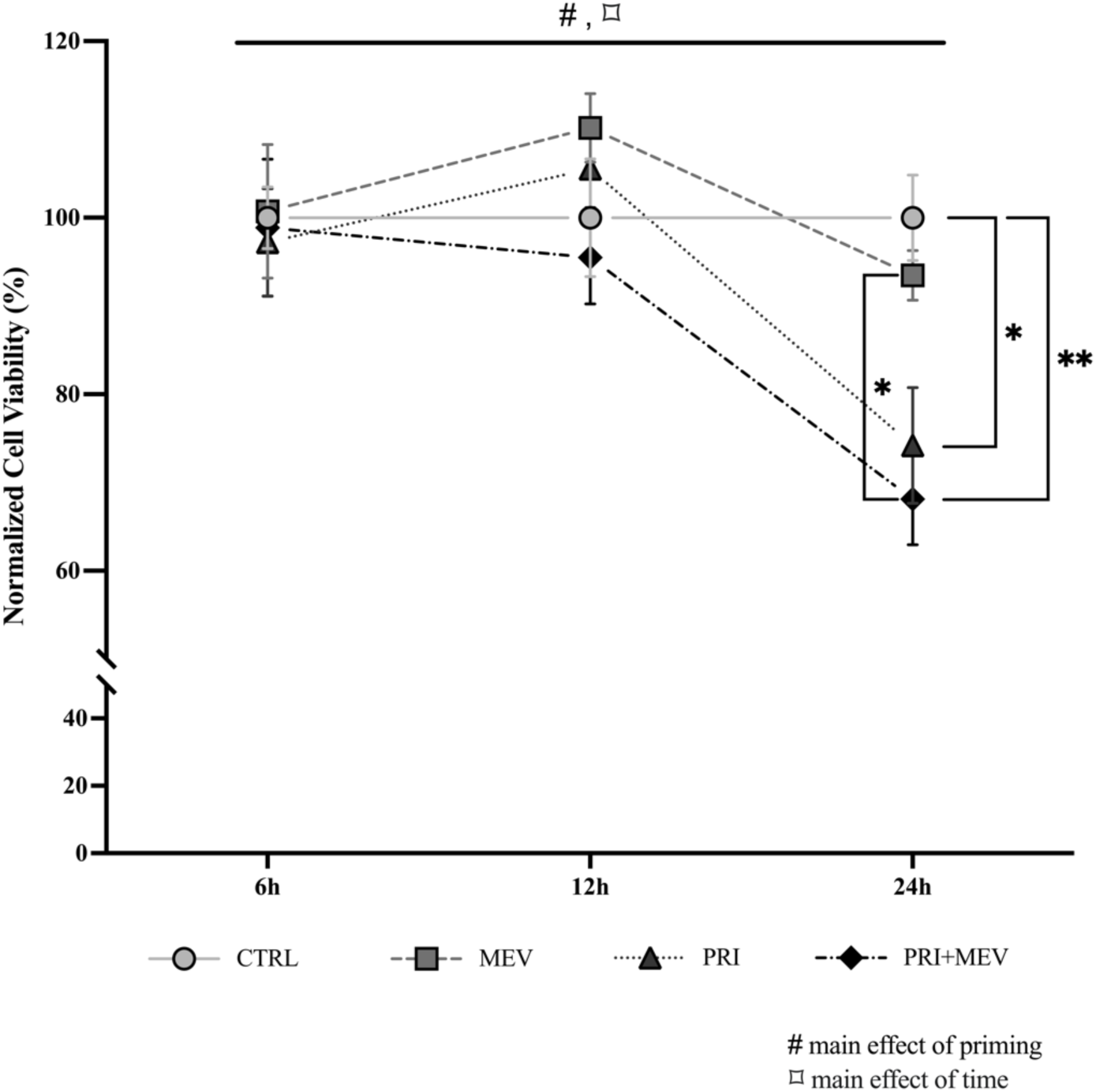
HMC3 cell viability with IFN-γ priming and MEV supplementation. HMC3 were plated at 2.5 x 10^4^ cells/well. At >90% confluence, a subset of cells was treated with 10 ng/mL IFN-γ (PRI and PRI+MEV groups). 24 hours post-IFN-γ treatment, a second subset of cells was supplemented with 200 µg MEV (MEV and PRI+MEV groups). A commercially available MTT (3-(4, 5-dimethylthiazolyl-2)-2, 5-diphenyltetrazolium bromide) assay was used to determine cell viability. The main effects of priming (denoted by ‘#’) and the main effects of time (denoted by ‘⌑’) are indicated. Significant differences between treatment groups were determined using a univariate general linear model with Tukey HSD (* p ≤ 0.05; ** p < 0.01; *** p < 0.001).

### 2.3 HSF1

Transcript abundance of HSF1 remained unchanged in homeostatic microglia at 6h (*t*(7) =-0.429, p = 0.681), 12h (*t*(7) =-4.10, p = 0.666), and 24h (*t*(5) = 0.927, p = 0.396) with MEV supplementation (**Figure 3a**). Similarly, in polarized microglia, no changes were seen in transcript at 6h (*t*(8) =-0.080, p = 0.938), 12h (*t*(8) =-1.478, p = 0.178), and 24h (*t*(8) =-1.498, p = 0.173) (**Figure 3b**).

**Figure 3.**
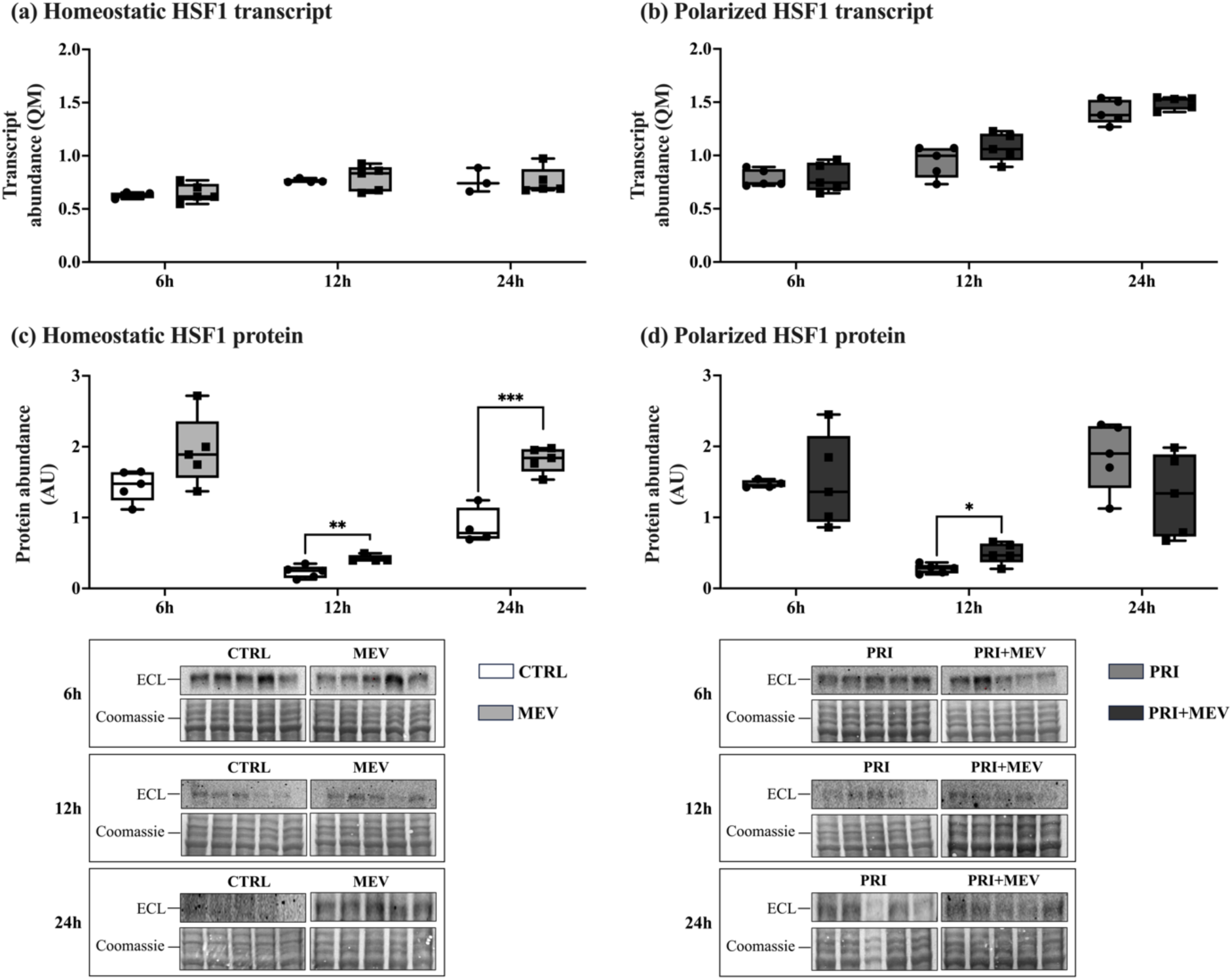
Transcript and protein abundance of HSF1 in response to IFN-γ priming and MEV supplementation, in homeostatic (CTRL and MEV) and polarized (PRI and PRI+MEV) HMC3 cells. **(a)** Homeostatic HSF1 transcript abundance. **(b)** Polarized HSF1 transcript abundance. **(c)** Homeostatic HSF1 protein abundance. **(d)** Polarized HSF1 protein abundance. RT-qPCR targets are normalized to the geometric mean of reference genes: *GAPDH* and *PKM*. Protein targets are normalized to total protein levels, with ECL and Coomassie-stained blot images displayed. Significant differences between treatment groups were determined using a two-tailed, independent samples t-test with Tukey HSD (* p ≤ 0.05; ** p < 0.01; *** p < 0.001).

At the protein level, in homeostatic microglia, HSF1 remained unchanged at 6h (*t*(8) =-2.055, p = 0.074), but increased with MEV supplementation at 12h (*t*(7) =-3.811, p = 0.007) and 24h (*t*(7) =-6.549, p < 0.001) (**Figure 3c**). Similarly, in polarized microglia, HSF1 abundance increased with MEV supplementation at 12h (*t*(8) =-2.991, p = 0.017), but remained unchanged at 6h (*t*(7) =-0.111, p = 0.906) and 24h (*t*(8) = 1.610, p = 0.146) (**Figure 3d**).

To examine HSF1 nuclear translocation (indicative of enhanced transcriptional activity), we assessed total HSF1 levels in the cytoplasm and the nucleus of HMC3 cells at 12h post-MEV supplementation. 12h post-supplementation was chosen for this analysis because the 12h timepoint had the most robust changes for HSF1 at the protein level in homeostatic and polarized microglia. In PRI+MEV cells, HSF1 was highly abundant in the nucleus (*F*(_3_,_16_) = 4.787, p = 0.014) compared to CTRL (Tukey HSD, p = 0.037), MEV (Tukey HSD, p = 0.025), and PRI (Tukey HSD, p = 0.037) cells (**Figure 4**). An increased abundance of HSF1 was also detected in the cytoplasm in PRI+MEV cells (*F*(_3_,_15_) = 10.929, p < 0.001) compared to CTRL (Tukey HSD, p < 0.001), MEV (Tukey HSD, p = 0.003) and PRI (Tukey HSD, p = 0.003) cells (**Figure 4**). The presence of histone H3, an integral component of the nucleosome, in the nuclear fraction and the absence in the cytoplasmic fraction confirms proper isolation of the cellular compartments.

**Figure 4.**
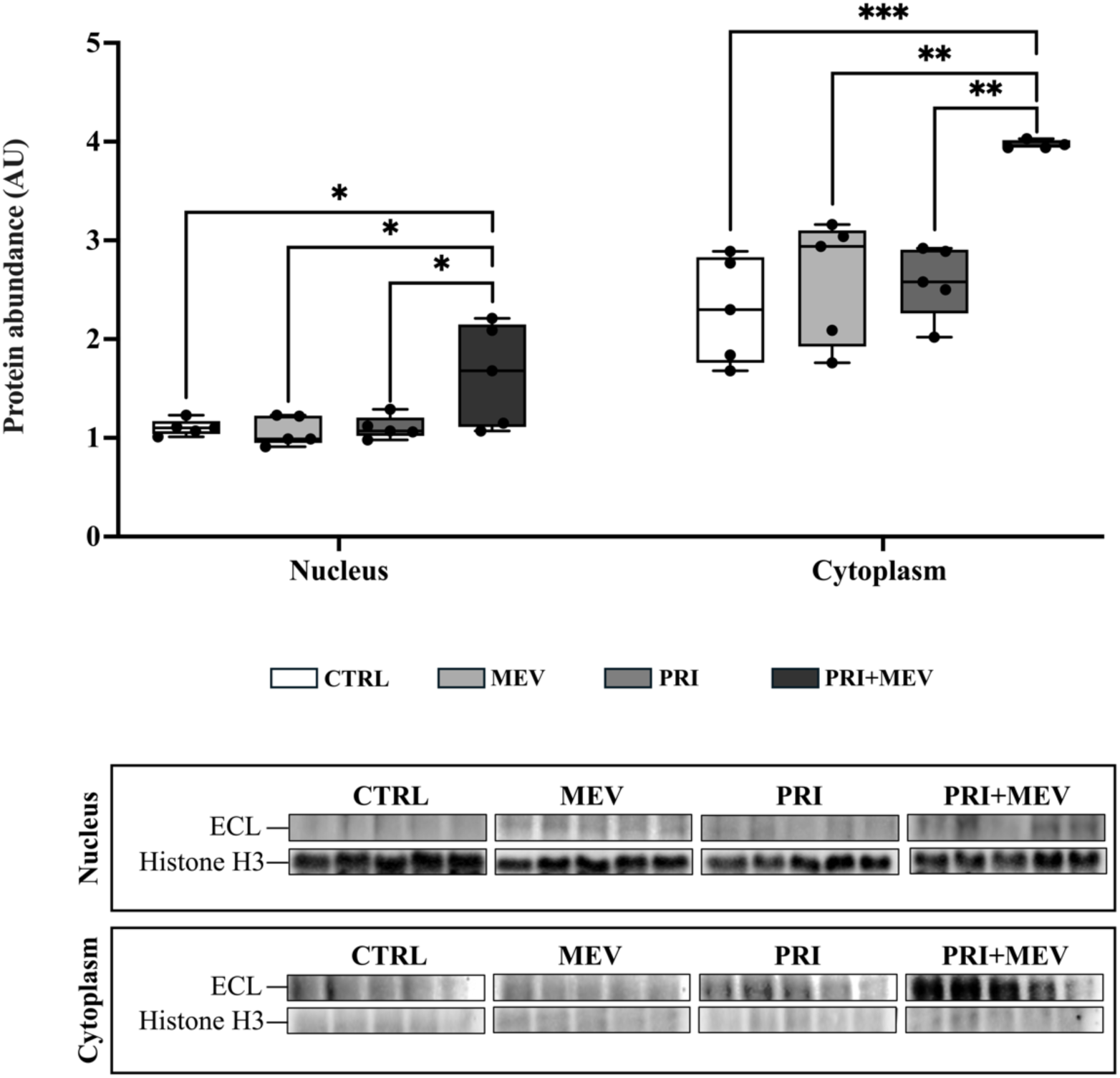
Cytonuclear HSF1 protein localization in response to IFN-γ priming and MEV supplementation, in the nuclear and cytoplasmic compartments of HMC3 cells. Protein targets are normalized to total protein levels, with ECL and Coomassie-stained blot images displayed. Significant differences between treatment groups were determined using a one-way ANOVA with Tukey HSD (* p ≤ 0.05; ** p < 0.01; *** p < 0.001).

### 2.4 Hsp70

In homeostatic cells, the transcript abundance of *HSPA1A* decreased with MEV supplementation at 6h (*t*(8) = 5.599, p < 0.001), but remained unchanged at 12h (*t*(8) =-0.433, p = 0.677) and 24h (*t*(7) =-1.726, p = 0.118) (**Figure 5a**). In polarized cells, there were no changes in transcript abundance of *HSPA1A* across the 6h (*t*(8) =-0.694, p = 0.507), 12h (*t*(8) = 0.726, p = 0.489), and 24h (*t*(8) =-0.217, p = 0.834) timepoints (**Figure 5b**).

**Figure 5.**
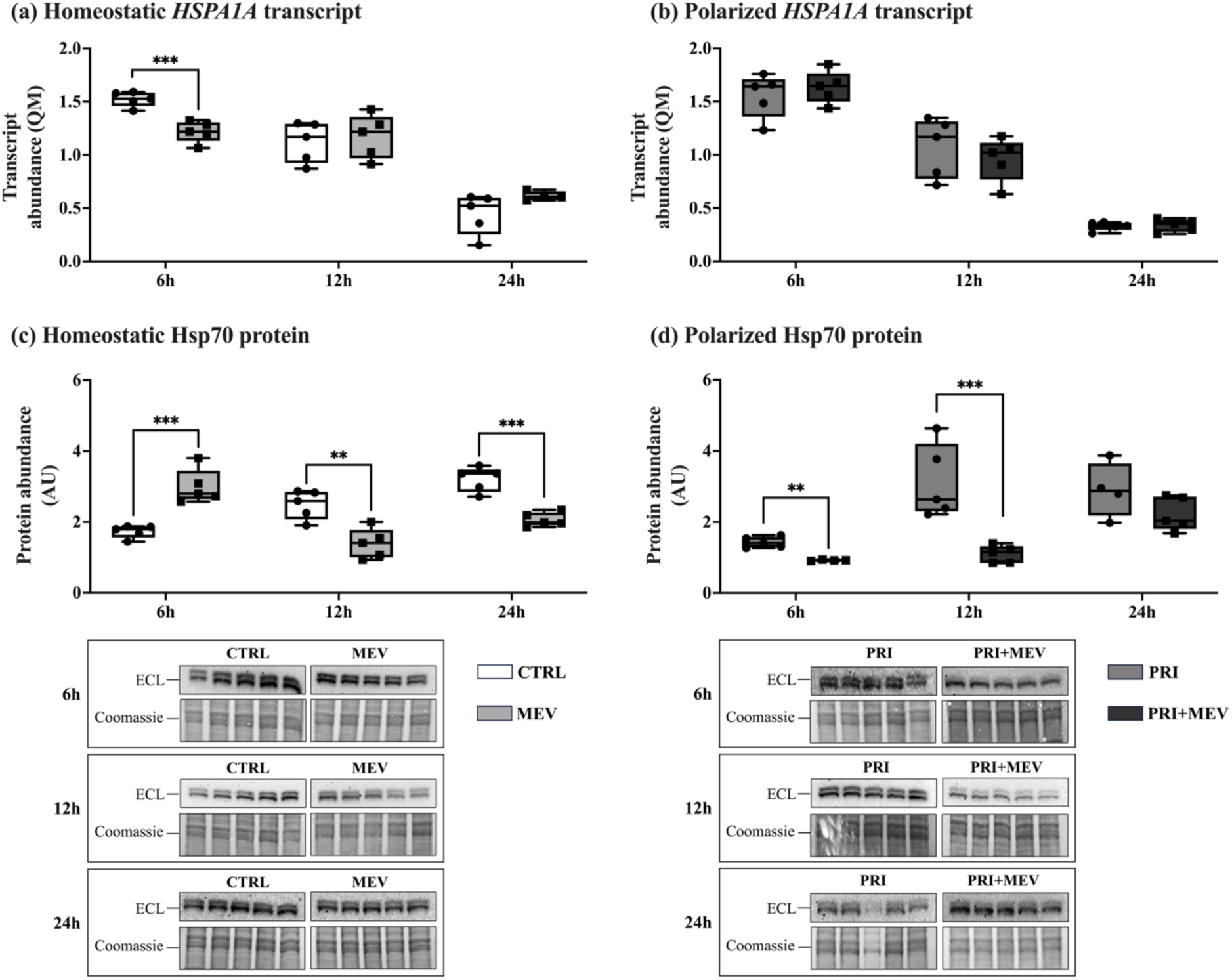
Transcript and protein abundance of Hsp70 (*HSPA1A*) in response to IFN-γ priming and MEV supplementation, in homeostatic (CTRL and MEV) and polarized (PRI and PRI+MEV) HMC3 cells. **(a)** Homeostatic *HSPA1A* transcript abundance. **(b)** Polarized *HSPA1A* transcript abundance. **(c)** Homeostatic Hsp70 protein abundance. **(d)** Polarized Hsp70 protein abundance. RT-qPCR targets are normalized to the geometric mean of reference genes: *GAPDH* and *PKM*. Protein targets are normalized to total protein levels, with ECL and Coomassie-stained blot images displayed. Significant differences between treatment groups were determined using a two-tailed, independent samples t-test with Tukey HSD (* p ≤ 0.05; ** p < 0.01; *** p < 0.001).

At the protein level, in homeostatic cells, Hsp70 levels increased at 6h (*t*(8) =-5.466, p < 0.001), but decreased at 12h (*t*(8) = 4.177, p = 0.003) and 24h (*t*(8) = 6.331, p < 0.001) (**Figure 5c**). In polarized cells, Hsp70 decreased at 6h (*t*(7) = 6.462, p = 0.002) and 12h (*t*(8) = 4.267, p = 0.010), but no change was observed at 24h (*t*(7) = 1.642, p = 0.145) (**Figure 5d**).

### 2.5 Hsp90

MEV supplementation did not change transcript abundance of *HSP90AA1* in homeostatic cells at 6h (*t*(8) =-0.211, p = 0.838), 12h (*t*(8) = 2.132, p = 0.066), and 24h (*t*(7) = 1.677, p = 0.128) (**Figure 6a**). Similarly, transcript abundance did not change in polarized cells at 6h (*t*(8) =-0.119, p = 0.908), 12h (*t*(8) =-1.225, p = 0.255), and 24h (*t*(7) =-1.695, p = 0.134) (**Figure 6b**).

**Figure 6.**
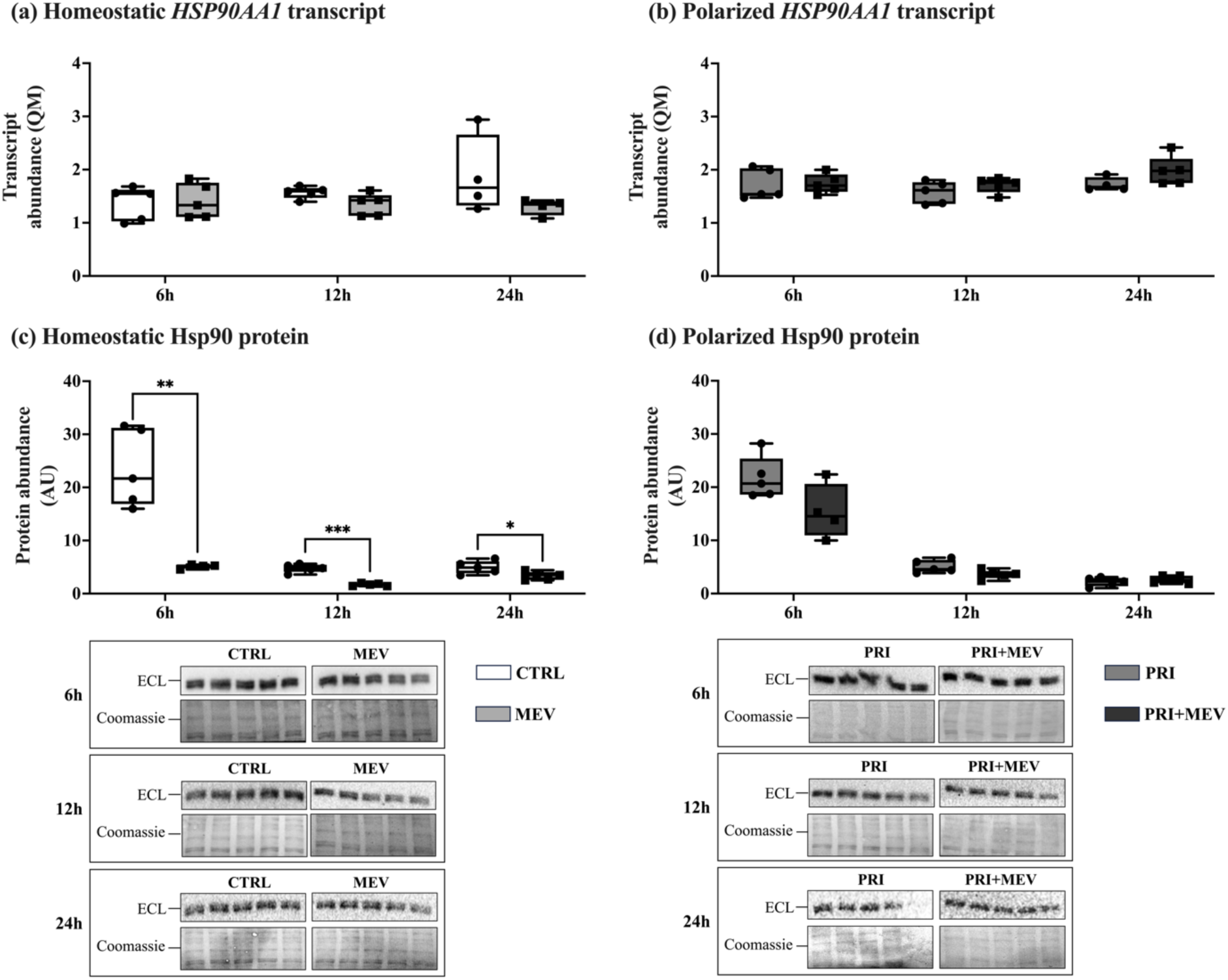
Transcript and protein abundance of Hsp90 (*HSP90AA1*) in response to IFN-γ priming and MEV supplementation, in homeostatic (CTRL and MEV) and polarized (PRI and PRI+MEV) HMC3 cells. **(a)** Homeostatic *HSP90AA1* transcript abundance. **(b)** Polarized *HSP90AA1* transcript abundance. **(c)** Homeostatic Hsp90 protein abundance. **(d)** Polarized Hsp90 protein abundance. RT-qPCR targets are normalized to the geometric mean of reference genes: *GAPDH* and *PKM*. Protein targets are normalized to total protein levels, with ECL and Coomassie-stained blot images displayed. Significant differences between treatment groups were determined using a two-tailed, independent samples t-test with Tukey HSD (* p ≤ 0.05; ** p < 0.01; *** p < 0.001).

At the protein level, MEV supplementation led to decreases in Hsp90 in homeostatic cells at the 6h (*t*(7) = 4.988, p = 0.005), 12h (*t*(8) = 8.526, p < 0.001), and 24h (*t*(8) = 2.469, p = 0.039) timepoints (**Figure 6c**). No changes were observed in polarized cells at 6h (*t*(7) = 2.091, p = 0.075), 12h (*t*(8) = 2.270, p = 0.053), or 24h (*t*(8) =-0.860, p = 0.415) (**Figure 6d**).

### 2.6 Hsp40

Transcript abundance of *DNAJB1* remained unchanged with MEV supplementation in homeostatic cells at 6h (*t*(8) = 0.304, p = 0.769), 12h (*t*(8) =-0.134, p = 0.897), and 24h (*t*(5) = 2.548, p = 0.051) (**Figure 7a**). Similarly, transcript abundance did not change in polarized cells at 6h (*t*(7) = 1.258, p = 0.224), 12h (*t*(8) =-0.918, p = 0.385), and 24h (*t*(7) =-1.798, p = 0.115) (**Figure 7b**).

**Figure 7.**
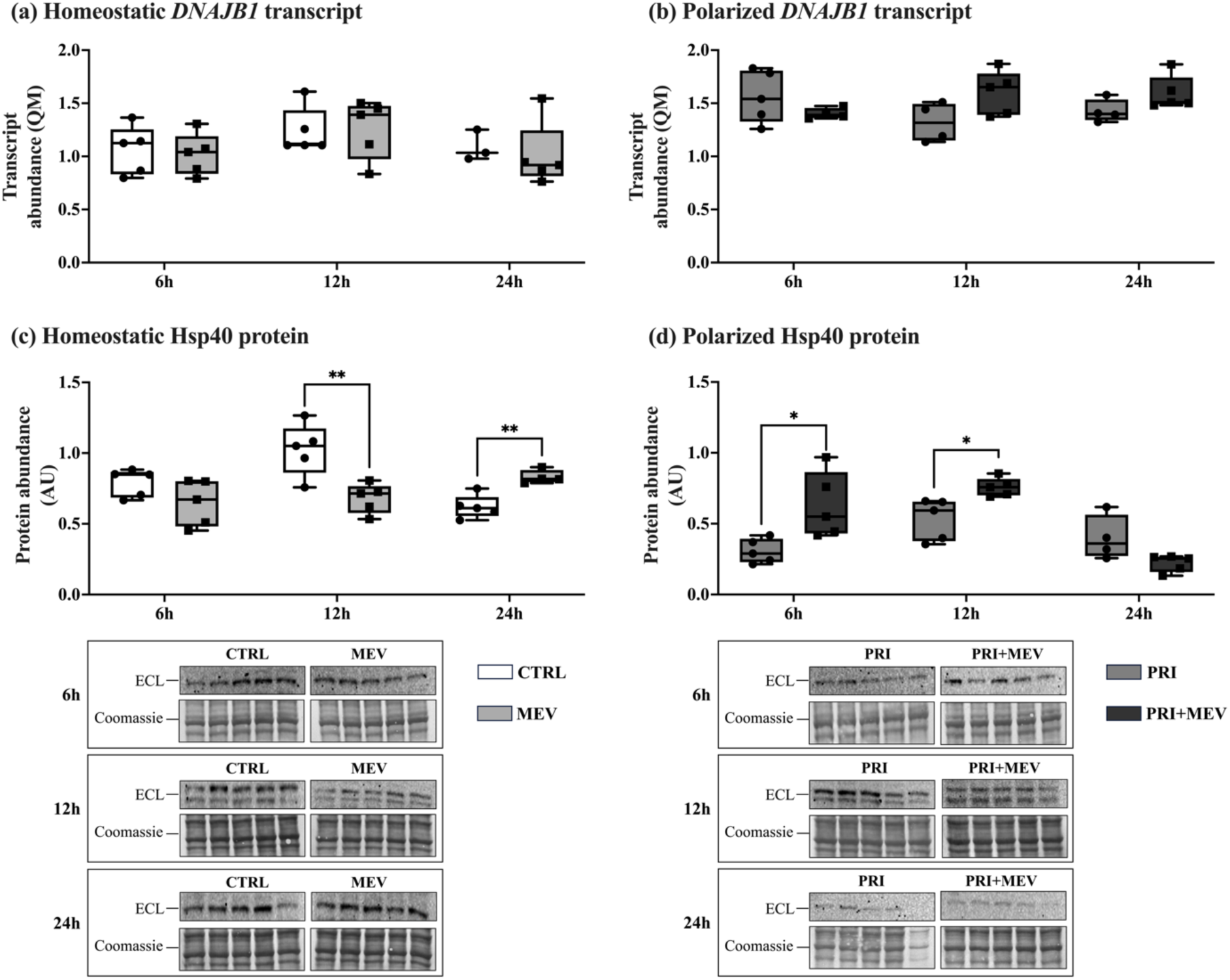
Transcript and protein abundance of Hsp40 (*DNAJB1*) in response to IFN-γ priming and MEV supplementation, in homeostatic (CTRL and MEV) and polarized (PRI and PRI+MEV) HMC3 cells. **(a)** Homeostatic *DNAJB1* transcript abundance. **(b)** Polarized *DNAJB1* transcript abundance. **(c)** Homeostatic Hsp40 protein abundance. **(d)** Polarized Hsp40 protein abundance. RT-qPCR targets are normalized to the geometric mean of reference genes: *GAPDH* and *PKM*. Protein targets are normalized to total protein levels, with ECL and Coomassie-stained blot images displayed. Significant differences between treatment groups were determined using a two-tailed, independent samples t-test with Tukey HSD (* p ≤ 0.05; ** p < 0.01; *** p < 0.001).

At the protein level, Hsp40 remained unchanged in homeostatic cells at 6h (*t*(8) = 1.689, p = 0.130), but we observed a decrease in abundance at 12h (*t*(8) = 3.616, p = 0.007), followed by an increase at 24h (*t*(7) =-4.497, p = 0.003) (**Figure 7c**). In polarized cells, Hsp40 increased at both 6h (*t*(8) =-2.894, p = 0.034) and 12h (*t*(8) =-3.188, p = 0.021), but remained unchanged at 24h (*t*(8) = 0.958, p = 0.366) (**Figure 7d**).

### 2.7 Hsp27

Transcript abundance of *HSPB1* remained unchanged with MEV supplementation in homeostatic cells at 6h (*t*(7) =-1.179, p = 0.277), 12h (*t*(8) = 1.591, p = 0.150), and 24h (*t*(7) =-0.002, p = 0.998) (**Figure 8a**). No changes were observed in polarized cells at 6h (*t*(7) = 0.959, p = 0.365), 12h (*t*(8) =-0.072, p = 0.945), or 24h (*t*(6) =-0.119, p = 0.909) as well (**Figure 8b**).

**Figure 8.**
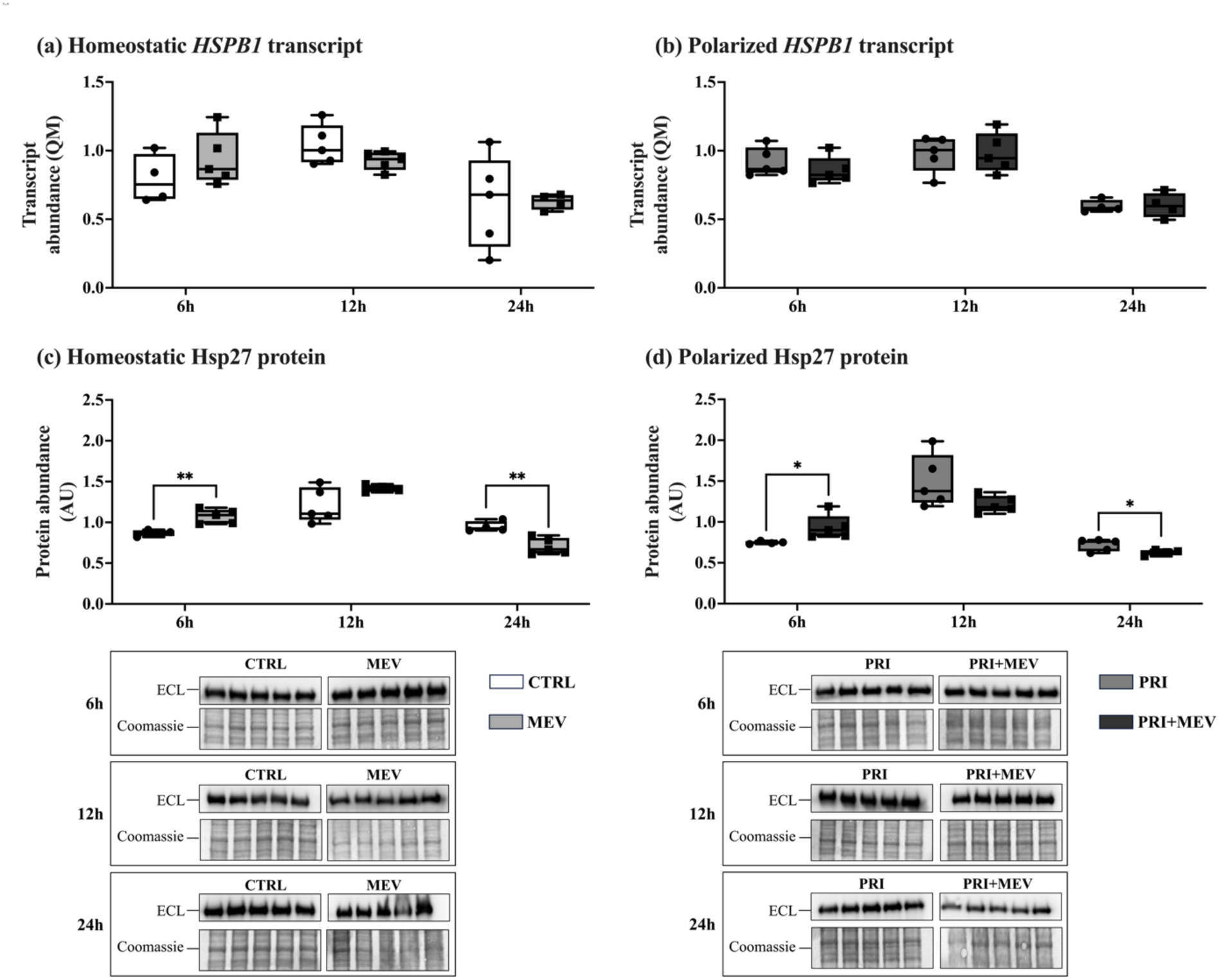
Transcript and protein abundance of Hsp27 (*HSPB1*) in response to IFN-γ priming and MEV supplementation, in homeostatic (CTRL and MEV) and polarized (PRI and PRI+MEV) HMC3 cells. **(a)** Homeostatic *HSPB1* transcript abundance. **(b)** Polarized *HSPB1* transcript abundance. **(c)** Homeostatic Hsp27 protein abundance. **(d)** Polarized Hsp27 protein abundance. RT-qPCR targets are normalized to the geometric mean of reference genes: *GAPDH* and *PKM*. Protein targets are normalized to total protein levels, with ECL and Coomassie-stained blot images displayed. Significant differences between treatment groups were determined using a two-tailed, independent samples t-test with Tukey HSD (* p ≤ 0.05; ** p < 0.01; *** p < 0.001).

Hsp27 protein abundance increased in homeostatic cells at 6h (*t*(7) =-4.293, p = 0.004), however, no change was observed at 12h (*t*(7) =-1.916, p = 0.091), and interestingly Hsp27 protein levels decreased at 24h (*t*(7) = 4.166, p = 0.004) (**Figure 8c**). In polarized cells, the same pattern was observed where Hsp27 protein levels increased with MEV supplementation at 6h (*t*(7) =-2.484, p = 0.042), with no change at 12h (*t*(8) = 1.853, p = 0.125), and decreased at 24h (*t*(7) = 2.270, p = 0.050) (**Figure 8d**).

### 3.0 Discussion

MEVs have immunomodulatory and anti-inflammatory characteristics. While MEVs have been shown to attenuate pro-inflammatory pathways and reverse proteotoxicity in peripheral tissues and organs, their bioactivity in the brain, in particular in immune-reactive macrophages, remains to be investigated. We investigated if and how MEVs may interact with the basal HSR, a principle pro-survival response, in homeostatic and IFN-γ polarized human microglia. Our overall findings suggest that MEV supplementation promotes the HSR, specifically by increasing HSF1 abundance and prolonging its activity, and increasing the abundance of candidate HSPs.

Extracellular vesicles (EVs) are complex molecules with diverse physiological and molecular roles that are produced and released by all cells [82]. This molecular and structural heterogeneity poses a challenge for researchers to define, isolate, and characterize EVs originating from different cells, tissues, and biological fluids [83]. Recommendations for characterization are thus published by the International Society for Extracellular Vesicles as the “Minimum Information for Studies of Extracellular Vesicles” (MISEV), which suggest that several aspects of EVs should be characterized for specificity, including but not limited to, quantification of particle size, morphology, and membrane protein composition [81]. We used three methods of characterization to satisfy the MISEV recommendations: NTA, TEM, and western immunoblotting. All three methods validated the isolation of MEVs and conferred to MISEV standards (**Figure 1**).

To ensure that IFN-γ priming and MEV supplementation did not impact cell viability and metabolic functions, we conducted an MTT assay. The cell viability remained unchanged at 6h and 12h in cells that received both treatments compared to control cells, suggesting that IFN-γ priming and MEV supplementation do not change the overall metabolic activity of the HMC3 cells (**Figure 2**). At 12h, MEV supplementation leads to a slight increase in cell viability (above 100%), compared to the control untreated cells. Live cells with an active metabolism use mitochondrial dehydrogenases to reduce MTT reagent into formazan, which can be analyzed as a proxy of cell viability [84]. Since formazan is insoluble and can accumulate as crystals inside the cell [84], likely the increase in absorbance with MEV supplementation that we are seeing at 12h is due to formazan crystal aggregation rather than a biologically relevant increase. As such, this increase should not be interpreted as an increase in cellular proliferation, viability, or mitochondrial output. Our results highlight some of the limitations of the MTT assay and the necessity for optimization and rational interpretation of results [85]. At 24h a significant priming effect is evident, where the PRI and PRI+MEV groups have reduced cell viability compared to the untreated control cells and homeostatic cells receiving MEVs. It is likely that while at 6h and 12h, MEVs were able to maintain a homeostatic microglial environment, at 24h the effects of the MEVs begins to dissipate, allowing the priming effects to dominate and decrease cell viability. Contrary to our findings, a study by La Torre et al. (2022) [86] reported that administration of EVs (10 μg/mL) isolated from BV-2 microglia for 24h did not impact cell viability of control BV-2 cells, or BV-2 cells pre-treated for 24h with 1 µg/mL LPS. Tong et al. (2021) [87] also reported that 24h incubation with isolated bovine MEVs (30 to 480 μg/mL) and LPS (100 ng/mL) did not affect the cell viability of RAW264.7 macrophage-like cells. It is possible that the reason we see a decrease in cell viability at 24h is due to the unique responses of immortalized human microglia to stress, which can depend on the cell-type, culture age, developmental stage, and host organism, as well as the type of stimulus, dosage, and duration [15].

HSF1 is the primary transcription factor that regulates the expression of HSP chaperones in response to and in preparation of proteotoxic stress [88]. Specifically, HSF1 can translocate to the nucleus and activate the transcription of candidate HSP genes within seconds to minutes of exposure to a proteotoxic stress or a cytokine trigger [64, 89]. Similarly, *HSF1* is typically expressed very early during cellular stress and is cell and time specific. Thus, HSF1 transcriptional output can be difficult to capture [90]. Correspondingly, at the transcript level, no significant changes are observed in homeostatic and polarized microglia at 6h, 12h, and 24h post-MEV supplementation (**Figure 3a-b**). Given that an immediate transcriptional response is required upon initiation of the HSR, HSF1 activation is primarily regulated at the protein level [64]. As a result, protein levels tend to change more drastically than messenger RNA (mRNA) levels, especially in experimental conditions with low temporal resolution, potentially resulting in discrepancies between mRNA and protein abundance data [90]. This is largely due to dynamic post-transcriptional (microRNAs; miRNAs) and post-translational regulatory mechanisms that shape gene expression patterns. Similar responses are also shown with other transcription factors that modulate pro-survival pathways, where transcriptional activation is induced rapidly within seconds to minutes of proteotoxic stress; for example, the hypoxia inducible factor-1 and transcription factors regulated by the Ras/Raf/MEK/ERK signaling pathway, such as NFκB, activator protein-1, cellular myelocytomatosis oncogene, and protein C-ets-1 [91–93]. The fact that the earliest timepoint we analyzed in our study was 6h post-MEV supplementation, it is likely that we had missed the transcriptional regulation but were able to capture changes in protein regulation. At the protein level, HSF1 increases in abundance in homeostatic microglia at 12h and 24h and increases in polarized microglia at 12h (**Figure 3c-d**). Cytonuclear translocation analysis of HSF1 shows that HSF1 abundance in the cytoplasmic and nuclear compartments is highest in polarized cells receiving MEVs (**Figure 4a-b**), compared to homeostatic cells. This suggests that HSF1 is not only increasing in polarized microglia, but that MEV supplementation in polarized cells leads to higher levels of active HSF1 in the nucleus. Taken together, these data suggest that increased HSF1 abundance with MEV supplementation implies an increase in HSR activation, and an overall increase in cytoprotective, pro-survival responses. Liao et al. (2023) [94] report that in a murine model of Parkinson’s disease (PD), overexpression of HSF1 represses microglia polarization, causing an increase in anti-inflammatory markers (TGF1β and IL-10), and a decrease in pro-inflammatory markers (IL-1β, IL-6, and TNFα), when compared to PD-induced microglia. Furthermore, Janus et al. (2022) [95] report that HSF1 can attenuate pro-inflammation by inhibiting candidate genes involved in the NFκB pathway, including NFκB repressing factor, NFκB inhibitor delta, TNF receptor associated factor 2 and TNF receptor associated factor 3 in human MCF7 ERα-positive breast cancer cells.

Similar to HSF1, the induction of HSP chaperones is primarily regulated at the protein level [64]. The Hsp70 chaperone plays an important role in protein folding and disaggregation and is an integral component of the Hsp70-Hop-Hsp90 complex [82, 96]. We observe a decrease in *HSPA1A* transcript levels in homeostatic microglia at 6h (**Figure 5a**), while *HSPA1A* transcript levels remained unchanged across other time points and in polarized microglia (**Figure 5a-b**). A potential explanation for this decrease in *HSPA1A* transcript at 6h is that miRNA cargo may induce post-transcriptional regulation of Hsp70 post-MEV supplementation. Several studies suggest that miRNA are the most likely candidates within MEV cargo to induce post-transcriptional regulation in recipient cells and systems [38–40]. However, the miRNA-induced post-transcriptional regulation that occurs in the brain requires further study, especially with regards to the interaction between MEV miRNA cargo and individual HSP chaperones. Among the ten most abundant MEV miRNA, several have been reported to downregulate *HSPA* family genes, including hsa-miR-30d-5p, hsa-miR-146b-5p, hsa-miR-21-5p, hsa-miR-30b-5p, and hsa-miR-30a-5p [97–99]. However, these studies did not look at these miRNAs from an MEV source or in the same cell line or pathophysiological condition, and thus would require further experiments to determine whether they play a role in regulating *HSPA1A*. The decrease in *HSPA1A* transcript at 6h in the homeostatic microglia are consistent with the Hsp70 protein levels at 12h and 24h (**Figure 5c**). Baseline homeostatic responses are typically activated early at the gene level, and these changes subsequently appear at the protein level several minutes to hours later. Further, as an integral part of the Hsp70-Hop-Hsp90 complex, Hsp70 plays a role in the negative regulation of HSF1 activation [64]. In particular, Hsp70 is responsible for interfering with the binding and stabilization of HSF1 monomers, preventing trimerization, and thus activation of HSF1 [64]. This negative feedback loop is designed to limit the hyperactivation and prolonged function of HSF1. At 12h and 24h, while Hsp70 protein levels decrease, HSF1 protein abundance increases in the same cells (**Figure 3a**). Likely, Hsp70 downregulation is resulting in the upregulation of HSF1 via the action of MEVs and their cargo. Interestingly, at 6h in homeostatic cells, Hsp70 protein abundance increases, unlike the decrease observed at 12h and 24h (**Figure 5c**). Hsp70 is highly enriched in MEVs [100–102]. Likely the increase being observed here results from the addition of Hsp70 from the MEV cargo to the Hsp70 protein endogenously produced by human microglia, rather than an increase in Hsp70 protein translation in response to the treatments. In polarized cells, we see a decrease in Hsp70 abundance at 6h and 12h, similar to the homeostatic 12h and 24h cells (**Figure 5d**). Once again, the decrease in Hsp70 abundance in polarized cells complements the increase in HSF1. This suggests that MEVs may be potentiating and prolonging HSF1 activity by reducing the amount of Hsp70 available in the cytoplasm to trimerize with Hop and HSP90, and thereby reducing the amount of Hsp70 binding to HSF1, preventing its nuclear translocation. Interestingly, although not statistically tested, we observed a gradual decrease in *HSPA1A* transcript levels with increasing time (from 6h to 24h) in polarized microglia (**Figure 5b**), which is the inverse of HSF1 transcript levels in polarized microglia which increase over time (**Figure 3b**). These observations suggest a time-related aspect of Hsp70-HSF1 negative regulation.

The Hsp90 chaperone is another key regulator of proteostasis and is a member of the Hsp70-Hop-Hsp90 complex [103]. No changes were seen in Hsp90 transcript abundance in homeostatic or polarized microglia (**Figure 6a-b**). However, at the protein level, Hsp90 abundance decreases across all timepoints in homeostatic microglia, and displays a decreasing trend in polarized microglia, although these results are not statistically significant (**Figure 6c-d**). These results follow the same pattern as Hsp70. As a binding partner within the Hsp70-Hop-Hsp90 complex, Hsp90 is involved in the disassembly of HSF1 trimers, and attenuates HSF1-mediated gene expression of candidate HSPs [64]. Our results show that with MEV supplementation, the same decreasing pattern observed in Hsp70 is also observed in Hsp90, once again with a corresponding increase in HSF1. These results suggest that the addition of MEVs results in the downregulation of Hsp70 and Hsp90 to upregulate HSF1 and maintain its activity over a longer period. Among the ten most abundant MEV miRNA, there are also several potential candidates involved in the downregulation of *HSPC* family genes that may be contributing to the responses we observe. These candidates include: hsa-miR-148a-3p, hsa-miR-200a-3p, hsa-miR-let-7b-5p, hsa-miR-146b-5p, hsa-miR-7f-5p, hsa-miR-21-5p, and hsa-miR-30a-5p [98, 99, 104–106]. Again, further experiments are required to investigate the potential role of these miRNAs in *HSP90AA1* post-transcriptional regulation.

Hsp40 is a co-chaperone that identifies and delivers misfolded proteins to its binding partner, Hsp70 for refolding by the Hsp70-Hop-Hsp90 complex [107]. However, we did not see significant differences in Hsp40 transcript levels in baseline or polarized microglia across timepoints (**Figure 7a-b**). At the protein level, in homeostatic cells, Hsp40 abundance decreases at 12h (**Figure 7c**). This follows the same pattern as homeostatic Hsp70 abundance at 12h. Typically, Hsp40 and Hsp70 follow the same regulatory patterns because Hsp40 enhances the ATPase activity of Hsp70 [108, 109]. Interestingly, we did not see a positive correlation between Hsp70 and Hsp40 protein levels at 24h in homeostatic cells, nor at 6h and 12h in polarized cells. At these timepoints, Hsp40 protein abundance increases with MEV supplementation (**Figure 7c-d**). Although the reason underlying this inverse regulation of Hsp70 and Hsp40 remains to be characterized, our study is not the first to report an inverse correlation. Chand et al. (2021) [110] report that during HIV-1 infection, Hsp70 (*HSPA1A*) was found to be downregulated, whereas Hsp40 (*DNAJB1*) was upregulated. This suggests that the protein association between Hsp70 and Hsp40 may be cell, tissue, and stress specific. As a possible explanation, we postulate that instead of increasing the ATPase activity of Hsp70, Hsp40 may be interacting with heat shock cognate 70 (Hsc70) [111, 112]. Hsc70 is also a part of the *HSPA* family (encoded by *HSPA8*) although it is constitutively expressed and is localized primarily in the cytoplasm [113]. Hsc70 is the only known chaperone involved in chaperone mediated autophagy (CMA), where misfolded clients are brought to the lysosome for degradation [114]. Hsp40 has been reported to stimulate Hsc70 ATPase activity, playing a critical role in Hsc70 substrate binding, and thus in preventing protein aggregation via CMA [112]. Further research is required to determine the specific Hsp40 binding partners in homeostatic and polarized microglia with MEV-supplementation.

Hsp27 is an ATP-independent chaperone that is integral in the rapid disaggregation of nuclear proteins [59, 78]. No changes are observed in Hsp27 transcript level in homeostatic or polarized microglia (**Figure 8a-b**). At the protein level, Hsp27 increases in both homeostatic and polarized cells at 6h, with no changes at 12h, but decreases at 24h (**Figure 8c-d**). In the absence of ATP, Hsp27 acts as a “holdase” and helps solubilize misfolded proteins, remodeling them into intermediate forms to be refolded by Hsp70 or Hsp90, preventing them from forming aggregates until the ATP-dependent chaperones are available [79, 80]. Given the temporal importance in Hsp27 activation as an ATP-independent chaperone, we postulate that its upregulation during the early stages of proteotoxic stress are critical, however, once other ATP-dependent chaperones are established, Hsp27-mediated protein rescuing becomes less crucial for maintaining homeostasis. However, further studies are required to delineate the mechanisms of Hsp27 regulation with MEV supplementation.

To our knowledge, our study is the first to explore the relationship between MEVs and the HSR, within the context of immune activation in a human macrophage, the microglia. Our findings suggest that MEV supplementation enhances the HSR, specifically by increasing HSF1 abundance and causing downregulation of its negative suppressors, Hsp70 and Hsp90 in homeostatic and polarized human microglia, suggesting a pro-survival role for MEVs in the brain. However, our study does not explore the mechanisms behind how MEV cargo may regulate the HSR. While additional components of MEV cargo may also contribute to HSR regulation (*e.g.*, short-chain fatty acids, other proteins), it is likely that miRNAs play a significant regulatory role. Future studies should explore specific miRNAs that are enriched in MEVs that also interact with and induce the downregulation of *HSPC* (*e.g.*, *HSP90AA1*) and *HSPA* family genes (*e.g., HSPA1A*). Among the top five most enriched MEV miRNA species, hsa-miR-148a-3p, hsa-miR-30d-5p, hsa-miR-200a-3p, and hsa-miR-let-7b-5p [115], *HSPA* genes have been previously reported to be downregulated specifically by hsa-miR-30d-5p [97, 98], and *HSPC* genes have been reported to be downregulated by hsa-miR-148a-3p, hsa-miR-200a-3p, and hsa-miR-let-7b-5p [98, 104, 105].

Although these studies were not necessarily conducted in the same cell type, pathophysiological condition, or in the context of MEV supplementation, it is a promising avenue for future research.

### 4.0 Conclusion

The cytoprotective and anti-inflammatory properties of MEVs have been largely established in peripheral tissues and organs *in vitro* and *in vivo*. MEVs are also emerging as candidates in translational medicine and nutritional science, owing to their therapeutic and bioengineering potential. However, despite these advances, the cytoprotective potential of MEVs in the CNS, particularly in brain macrophages, remains to be studied. Our study is the first to investigate the interactions between MEVs and pro-survival pathways in human microglia, in the context of alleviating neuroinflammation. We found that MEV supplementation promotes the HSR in homeostatic and polarized human microglia. Specifically, MEV supplementation increases the abundance of HSF1, the main transcription factor, while downregulating Hsp70-Hop-Hsp90 complex formation. Since Hsp70 and Hsp90 are implicated in the negative regulation of HSF1, their downregulation with MEV supplementation allows for prolonged HSR activation, exerting robust and continued pro-survival benefits in response to IFN-γ induced polarization in microglia. While further research needs to be conducted to investigate which component of the MEV cargo are inducing these effects, our results support the potential of MEVs to be used as a biotherapeutic to alleviate neuroinflammation.

### 5.0 Materials and Methods

### 5.1. MEV isolation and characterization

Unpasteurized human donor milk (n = 11 anonymous donors) was obtained from NorthernStar Mothers Milk Bank (Calgary, AB, CA) and pooled to create a homogeneous mixture. The milk was pooled to limit individual variability in milk composition of the donors (*e.g.*, metabolic/health status, milk stage). The milk was serially centrifuged to remove creams, fats, and milk fat globular membranes (twice at 3,000 x g, 10 mins, 22 °C), cellular debris (twice at 1,2000 x g, 10 mins, 4 °C), and milk cells (twice at 21,500 x g, 30 mins, 4 °C, followed by once at 21,500 x g, 60 mins, 4 °C). The remaining whey fraction was filtered using a 0.45 µm syringe filter (UltiDent: 229749) to remove residual milk cells and debris. Casein protein was removed using acetic acid precipitation (1:1000, v/v), and the whey supernatant was centrifuged (4,500 x g, 30 mins, 4 °C) and filtered through a 0.22 µm syringe filter (UltiDent: 229747) to remove casein proteins [116]. To isolate the MEVs, the supernatant was ultracentrifuged at 100,000 x g for 120 mins at 4 °C (Beckman-Coulter XL-100; SW55TI Swing-Bucket Rotor; no brake). The MEV pellet was washed in filtered 1X phosphate buffered saline (PBS) (1:1, v/v) and a second ultracentrifugation was completed at the same settings. The MEV pellet was resuspended in filtered 1X PBS (1:2, v/v to original whey volume) to create an enriched MEV sample used for supplementation experiments, and the supernatant from the second ultracentrifugation step was used as an EV-depleted negative control [31].

Following isolation, MEVs were characterized in accordance with the MISEV 2023 guidelines [81]. MEV particle size and concentration, and negative controls were quantified using NTA (Malvern Instruments Ltd.; NanoSight NS300) at the Hospital for Sick Children’s Structural and Biophysical Core Facility (Toronto, ON, CA). MEVs were diluted (1:300, v/v in 1X filtered PBS) [31] using an sCMOS camera (level 15, detection threshold 10) using a 532nm green laser (3x replicates at 30s capture speeds) as per Wijenayake et al., (2021) [31].

MEV morphology and integrity was characterized using TEM (FEI Talos F200x S/TEM) as per Wijenayake et al., (2021) [31] at the Manitoba Institute of Materials (Winnipeg, MB, CA). MEVs were loaded onto 400 mesh carbon-coated formvar film copper grids (Electron Microscopy Sciences: CF400-CU-50) suspended in 1X PBS (1:3, v/v), and negatively stained using 2% uranyl acetate.

The presence of three endosome-specific proteins, including two tetraspanins (CD9, CD81) and an EV biogenesis factor (syntenin-1), were detected using western immunoblotting. Protein abundance of calnexin, a cellular marker of endoplasmic reticulum, was used as the negative control to identify cellular protein contamination in the MEV isolates. Total soluble protein isolated from HMC3 cells was used as a cellular control. Total soluble protein was extracted by incubating MEVs and HMC3 cells in cell extraction buffer (Invitrogen: FNN0011), protease inhibitor cocktail (1:20, v/v) (BioShop: PIC001.1) and phenylmethanesulfonyl fluoride (1 mM) (BioShop: PMS123.5) for 30 minutes on ice with intermittent vortexing, followed by centrifugation (13,000 rpm, 10 min, 4 °C). Protein concentrations were quantified via a bicinchoninic acid (BCA) assay using the Pierce^™^ BCA Protein Assay kit (Thermo Scientific: 23227), according to the manufacturer’s instructions. Absorbance readings were obtained at 562 nm using a BioTek Synergy H1 Multimode microplate reader (Agilent: BTSH1M2SI). Bromophenol blue loading dye containing sodium dodecylsulfate (SDS) (Bioshop: SDS001.1) and *β-*mercaptoethanol (10%, v/v) (Bioshop: MERC002.500) was added to the samples, vortexed, and heated to 95 °C for 10 minutes to denature the proteins. Converted lysates were stored at-20 °C. Western immunoblotting parameters for MEV protein biomarkers are listed in **Table S1**. Antibody information for MEV biomarkers is listed in **Table S2**. Enhanced chemiluminescence (ECL) and Coomassie images for MEV protein biomarkers can be found in **Figure S1.** Isolated MEVs were stored at-80 °C for supplementation experiments.

### 5.2 Human microglia clone 3 (HMC3) culture

Immortalized HMC3 cell line (ATCC; CRL-3304; lot number: 70026037) was donated by Dr. Patrick O. McGowan (1 vial; Passage (P) 1; 2.26 x 10^6^ cells/mL) from the University of Toronto. The transfer of the culture to Dr. Sanoji Wijenayake at The University of Winnipeg (received on dry ice, February 14, 2023) followed permit requirements and material transfer agreements from ATCC and the Canadian Food Inspection Agency (compliance letter ID: CL-2022-0019-4). Cells were cultured with Eagle’s Minimum Essential Media (EMEM) (ATCC: 30-2003) supplemented with 10% fetal bovine serum (FBS) (ATCC: 30-2020) and maintained in a humidified incubator at 37 °C with 5% CO_2_, as per manufacturer’s recommendations. Complete media was replaced every 48 hours to prevent excessive acidification [117]. 0.25% Trypsin with 0.52 nM EDTA (ATCC: 30-2101) was used for cell detachment and harvesting. A hemocytometer was used for manual cell counting. Specifically, live cells in each of four quadrants were manually counted, and cell density and total cell counts were determined using the following formulae:

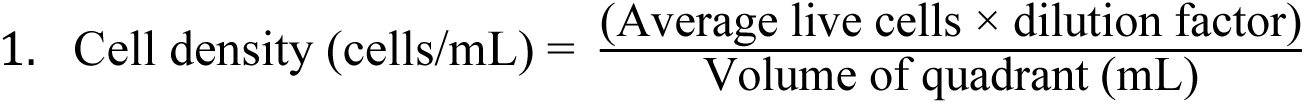

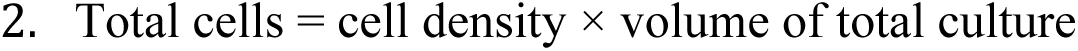

### 5.3 MEV supplementation

P8 cells were plated onto 6-well culture plates (Sarstedt: 83.3920) at 2 x 10^5^ cells/well (n = 5 biological replicates/treatment). P8 HMC3 cells have been used by previous *in vitro* studies and were reported to have stable transcriptional output [118].

A subset of cells was treated with 10 ng/mL IFN-γ (Millipore-Sigma: SRP3058) diluted in EMEM with 10% EV-depleted FBS and incubated for 24h (PRI and PRI+MEV cells). 10 ng/mL IFN-γ was selected to polarize microglia as per Franck and Storm *et al.* (2024) [15] and Peudenier et al. (1991) [119]. After 24h IFN-γ incubation, a second subset of cells were supplemented with 200 µg MEVs (MEV and PRI+MEV cells). All cells (CTRL, MEV, PRI and PRI+MEV) were harvested at 6h, 12h, and 24h post-MEV supplementation (**Figure 9)**. The 200 µg MEV dosage was selected based on published literature and has been shown to induce molecular responses in human and rodent cell lines [118, 120].

**Figure 9.**
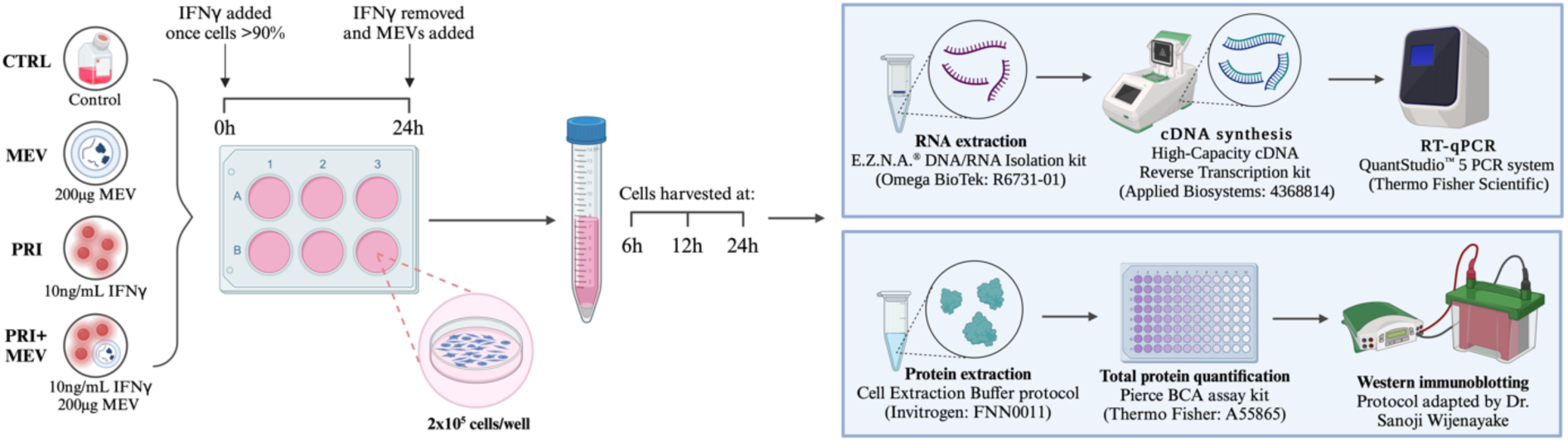
Schematic of microglia priming, supplementation, harvesting, and molecular analysis. HMC3 were plated at 2 x 10^5^ cells/well. At >90% confluence, a subset of cells was treated with 10 ng/mL IFN-γ (PRI and PRI+MEV groups). 24 hours post-IFN-γ treatment, a second subset of cells was supplemented with 200 µg MEV (MEV and PRI+MEV groups). All cells were harvested at 6h, 12h, and 24h. Total soluble RNA was extracted, and 3000 ng were reverse transcribed into cDNA. RT-qPCR was used to analyze transcript abundance of candidate heat shock response genes. Total soluble proteins were extracted, quantified using BCA assay, and converted to lysate using 2x SDS. Western immunoblotting was used to analyze protein abundance of candidate heat shock proteins.

EV-depleted FBS was used for MEV supplementation experiments to minimize the biological influence of FBS EVs in the experiment which may confound MEV effects [121]. FBS was ultracentrifuged at 100,000 x g for 18h at 4 °C (Beckman-Coulter XL-100; SW55TI Swing-Bucket Rotor; no brake) to pellet FBS EVs. The pellet was discarded, and the supernatant was filtered through a 0.22 µm syringe filter (UltiDent: 229747) prior to use in complete media.

### 5.4 Cell viability and mitochondrial output

A CyQUANT MTT Cell Viability Assay kit (Invitrogen: V-13154) was used to quantify the mitochondrial output of HMC3 with IFN-γ treatment and MEV supplementation as per the manufacturer’s instructions. Absorbance readings were obtained at 570 nm using a BioTek Synergy H1 Multimode microplate reader (Agilent: BTSH1M2SI). Cells were treated with 10 ng/mL IFN-γ for 24h and/or 200 µg MEVs, with untreated HMC3 cells used as a baseline control, and harvested at 6h, 12h and 24h, as previously described. Based on previous experiments by Franck and Storm *et al*. (2024) [15], 2.5 x 10^4^ cells/well was selected as the optimal plating density and was used for MTT cell viability assays (n = 5 biological replicates/treatment). Cell viability was calculated by normalizing blank-adjusted absorbance values to the average of the control cells [122, 123].

### 5.5 RNA extractions and cDNA synthesis

Total soluble RNA (≥18 nucleotides) was extracted from HMC3 cells using the E.Z.N.A.^®^ DNA/RNA Isolation kit (Omega BioTek: R6731-01) according to the manufacturer’s instructions. RNA concentration (ng/µL) and purity (A260:280 and A260:230 ratios) were measured using a NanoDrop One/One^C^ Microvolume-UV/Vis spectrophotometer (ThermoFisher Scientific: ND-ONE-W). Samples with A260:A280 ratios of 1.8-2.0 were used for complementary DNA (cDNA) synthesis. An RNA Clean & Concentrator kit (Zymo Research: R1017) was used to clean select RNA samples with A260:280 ratios <1.8-2.0, according to the manufacturer’s instructions. RNA stability and integrity (**Figure S2-S4**) were verified by resolving samples on a 1% TAE-agarose gel with 2x RNA loading dye (1:1, v/v) (Life Technologies: R0641) stained with Red Safe dye (FroggaBio: 21141) using a Sub-Cell GT Agarose Gel Electrophoresis System (BioRad:1704401). 3000 ng of total soluble RNA was reverse transcribed into cDNA using the High-Capacity cDNA Reverse Transcription kit (Applied Biosystems: 4368814) according to the manufacturer’s instructions. A T100™ Thermal Cycler (Bio Rad) was used for synthesis using the following amplification parameters: 25 °C for 10 mins; 37 °C for 120 mins; 85 °C for 5 mins; hold at 4 °C. RNA and cDNA were stored at-80°C.

### 5.6 Quantitative reverse transcription polymerase chain reaction (RT-qPCR)

mRNA-specific primers were designed using nucleotide sequence information available at the National Center for Biotechnology information (NCBI): http://www.ncbi.nlm.nih.gov or obtained from previously published literature. Primer pairs were assessed for compliance with the Minimum Information for Publication of Quantitative Real-Time PCR Experiments (MIQE) guidelines [124] using sequence information available on NCBI as well as using the OligoAnalyzer^™^ Tool from Integrated DNA Technologies (IDT, Coralville, Iowa, USA). Primer parameters are listed in **Table S3.** Primer bioinformatics are listed in **Table S4**.

RT-qPCR was performed using a QuantStudio^™^ 5 PCR system (Applied Biosystems: 96- well and 0.2 mL block) using the Fast SYBR^™^ Green Master Mix chemistry (Applied Biosystems: 4385612). Annealing temperatures for primer pairs were determined by testing a range of temperatures approximately ± 5 °C of the melting temperature of the forward primer (5’-3’), using the Veriflex setting of the QuantStudio^™^ 5 PCR system. Temperature testing was conducted using a pool sample (10 ng/µL) representing all test samples. Melt curve analysis was conducted to determine the specificity of the primer pairs. Primer pairs that generated a single, sharp peak void of primer dimers and a derivative reporter >200,000 were used for quantification. A 9-point standard curve ranging from 1000 ng/μL to 3.91 ng/μL was used to determine the optimal cDNA amount to load per primer pair. Analyses of all samples were done in triplicate and an inter-plate converter (IPC) was used on all plates to account for variability between plates and runs. Absolute transcript abundance of HSF1 and candidate HSPs (Hsp70, Hsp90, Hsp40, Hsp27) were analyzed using quantity means (QMs). The internal controls GAPDH, PKM, and 18S were selected as reference genes. Normfinder software (RRID: SCR_003387) [125] was used to confirm the thermal stability and intra/intergroup variability of the reference genes in response to IFN-γ treatment and MEV supplementation. All targets were normalized against the geometric mean of the three reference genes [126].

### 5.7 Total and cyto-nuclear protein extractions

Total soluble protein was extracted from frozen HMC3 pellets using cell extraction buffer with PIC, and PMSF according to the same protocols used for MEV protein extractions. Cyto-nuclear proteins were extracted using the NE-PER^™^ Nuclear and Cytoplasmic Extraction kit (Thermo Scientific: 78833) according to manufacturer’s instructions. Protein concentrations were quantified using a BCA assay. Absorbance readings were obtained at 562 nm. Lysates were solubilized using bromophenol blue loading dye with SDS and *β-*mercaptoethanol followed by mechanical denaturation (vortexing and temperature). Lysates were stored at-20 °C.

### 5.8 Western immunoblotting

6-15% SDS-polyacrylamide gel electrophoresis (SDS-PAGE) was used to resolve protein lysates according to target molecular weights. A 7-point standard curve using protein aliquots from all test samples (5 μg to 40 μg) was used to determine the amount of protein to load per target. BLUelf pre-stained protein ladder (FroggaBio: PM008-0500) was used as the molecular weight standard. After determining the optimal loading amount, samples were run with n = 5 biological replicates/treatment/timepoint. SDS-PAGE was conducted for 60-80 minutes in 1x Tris-Glycine running buffer (0.3%, w/v Tris-base; 14.4%, w/v glycine; 1%, w/v SDS) at 180V in a Sub-Cell GT Electrophoresis Cell (BioRad: 1704401). Proteins were transferred from the gel onto a 0.45 µm PVDF membrane (BioRad: 1620174) using a Trans-Blot Turbo Transfer System (BioRad: 1704150). PVDF membranes were blocked using 1-10% casein-TBST solution (30 minutes, 22°C) on a rocker and incubated with primary antibody (1:1000, v/v) followed by goat HRP-conjugated anti-rabbit IgG secondary antibody (1:5000, v/v or 1:10,000, v/v) on a rocker (45 minutes, 22°C). Blots were visualized using western ECL substrate solution (Thermo Scientific: 34580) and chemiluminescence imaging (ChemiDoc XRS+; BioRad: 1708265) with Image Lab Software (version 6.1). The immunoblots were stained with Coomassie brilliant blue (Bioshop: CBB555.10) solution (0.25%, w/v Coomassie blue salt; 7.5% acetic acid; 50%, v/v methanol) for 2 minutes and destained with destain solution (25%, v/v methanol; 10%, v/v acetic acid; ddH_2_O) for 5 minutes at room temperature. Coomassie-stained immunoblots were used for total protein normalization, where intensity of the target ECL bands was normalized against a set of proteins with constant abundance, not including the target protein. ImageJ (version 1.5.3) was used to quantify protein abundance [127]. Western immunoblotting parameters for HSR targets are listed in **Table S5**. Antibody information for HSR targets is listed in **Table S6**. ECL and Coomassie images for HSF1 and HSP targets are displayed in **Figure S5-S9**.

### 5.9 Statistical analysis

Statistical analysis was carried out using SPSS version 29.0.2.0 (IBM Corp.), and figures were constructed using GraphPad Prism version 9.5.1 and BioRender.com. Shapiro-Wilk tests were used to test for normality and Levene’s tests were used to assess homogeneity of variance. All data were normally distributed (p > 0.05) and independent by design. As such parametric testing was used. A two-tailed independent sample t-test was used to determine significant differences across homeostatic or polarized microglia with MEV supplementation (p ≤ 0.05). One-way analysis of variance (ANOVA) was used to determine significant differences across treatment groups for the cyto-nuclear HSF1 data (p ≤ 0.05). Univariate general linear models were used to determine significant differences across treatment groups for the MTT cell viability assay (p ≤ 0.05) to determine the main effect of priming, main effect of MEV supplementation and their associations. Extreme outliers with an interquartile range > 3 were identified using the SPSS boxplot outlier function and removed from the dataset, only when necessary. No more than one outlier was removed per treatment group. Pairwise comparisons between treatment groups were determined using Tukey HSD post-hoc (p ≤ 0.05).

## Abbreviations

ATP: Adenosine triphosphate
BCA: Bicinchoninic acid
cDNA: Complementary DNA
CMA: Chaperone mediated autophagy
CNS: Central nervous system
DNA: Deoxyribonucleic acid
DOHaD: Developmental Origins of Health and Disease
ECL: Enhanced chemiluminescence
EMEM: Eagle’s minimum essential media
EV: Extracellular vesicle
FBS: Fetal bovine serum
HMC3: Human microglia clone 3
HSF: Heat shock transcription factor
HSP: Heat shock protein
HSR: Heat shock protein response
IFN-γ: Interferon-gamma
IPC: Inter-plate converter
MEV: Milk-derived extracellular vesicle
miRNA: MicroRNA
MISEV: Minimum information for studies of extracellular vesicles
MIQE: Minimum Information for Publication of Quantitative Real-time PCR
mRNA: Experiments Messenger RNA
MTT: 3-[4,5-dimethylthiazol-2-yl]-2,5 diphenyl tetrazolium bromide
NCBI: National Center for Biotechnology Information
NFκB: Nuclear factor kappa B
NTA: Nanoparticle tracking analysis
PBS: Phosphate-buffered saline
PD: Parkinson’s disease
RNA: Ribonucleic acid
RT-qPCR: Quantitative reverse transcription polymerase chain reaction
SDS: Sodium dodecylsulfate
SDS-PAGE: Sodium dodecylsulfate polyacrylamide gel electrophoresis
TEM: Transmission electron microscopy
UTR: Untranslated region

## Data Availability Statement

The authors confirm that the data supporting the findings of this study are available within the article and its supplementary materials.

## Acknowledgements

We would like to thank Dr. Patrick O. McGowan (Ph.D.) from the University of Toronto for providing a vial of the HMC3 cell line for propagation and use in the Wijenayake Lab at The University of Winnipeg. We would like to thank the research technicians at the Structural and Biophysical Core Facility at the Hospital for Sick Children (Toronto, ON, Canada) for conducting nanoparticle tracking analysis of MEV samples. We would like to thank the support staff at the Manitoba Institute for Materials at the University of Manitoba (Winnipeg, MB, Canada) for sharing their facilities and use of the FEI Talos F200x S/TEM transmission electron microscope. We would also like to thank the support staff at The University of Winnipeg for their assistance.

## Author contributions

JAS: Data curation, formal analysis, investigation, validation, writing – original draft, writing – review & editing.

JL: Data curation.

MFO: Data curation.

SW: Conceptualization, formal analysis, funding acquisition, supervision, writing – original draft, writing – review & editing.

