## Supplementary Materials for "Human milk-derived extracellular vesicles promote the heat shock response in polarized microglia"

**Table S1. Western immunoblotting parameters for MEV biomarkers, CD9, CD81, syntenin-1, and the cellular protein, calnexin**. Targets are identified with the corresponding molecular weight (kDa), SDS-PAGE gel percentage (%), amount of protein loaded (µg), SDS-PAGE gel run time and voltage, transfer settings (time, voltage, amperage), casein blocking (percentage, time), primary antibody incubation parameters (concentration, time, temperature), and secondary antibody incubation parameters (concentration, time, temperature).

| Target | Molecular weight (kDa) | Gel % | Protein loaded (µg) | SDS-PAGE run | Transfer settings | Casein blocking | Primary antibody incubation | Secondary antibody incubation |
|---|---|---|---|---|---|---|---|---|
| Calnexin | 90 | 8 | 30 | 85 min, 180 V | 40 min, 25 V, 2.5 A | 1%, 30 min | 1:1000, 20h, 4 °C | 1:15,000, 45 min, 22 °C |
| CD9 | 22 | 15 | 30 | 80 min, 180 V | 7 min, 25 V, 2.5 A | 1%, 30 min | 1:1000, 20h, 4 °C | 1:15,000, 45 min, 22 °C |
| CD81 | 22, 24, 35 | 15 | 30 | 80 min, 180 V | 7 min, 25 V, 2.5 A | 2.5%, 30 min | 1:1000, 20h, 4 °C | 1:15,000, 45 min, 22 °C |
| Syntenin-1 | 30 | 10 | 30 | 80 min, 180 V | 20 min, 25 V, 1.5 A | 1%, 30 min | 1:1000, 20h, 4 °C | 1:15,000, 45 min, 22 °C |

**Table S2. Primary antibodies used for western immunoblotting of the MEV biomarkers**. Information includes the commercial antibody name, protein target, purchasing company and catalogue number, antigen species targeted, molecular weight, clonality, and host isotype. All antibodies recognize *Homo sapiens*.

| Antibody name | Target | Company | Catalogue number | Antigen species | Molecular weight (kDa) | Clonality | Host Isotype |
|---|---|---|---|---|---|---|---|
| Calnexin (C5C9) Rabbit mAb #2679 | Calnexin | Cell Signalling | #2679 | Human | 90 | Monoclonal | Rabbit IgG |
| CD9 (D8O1A) Rabbit mAb #13174 | CD9 | Cell Signalling | #13174 | Human | 22 | Monoclonal | Rabbit IgG |
| CD81 (D3N2D) Rabbit mAb #56039 | CD81 | Cell Signalling | #56039 | Human | 22, 24, 25 | Monoclonal | Rabbit IgG |
| Syntenin-1/MDA9 (E2I9L) Rabbit mAb #27964 | Syntenin-1 | Cell Signalling | #27964 | Human | 30 | Monoclonal | Rabbit IgG |

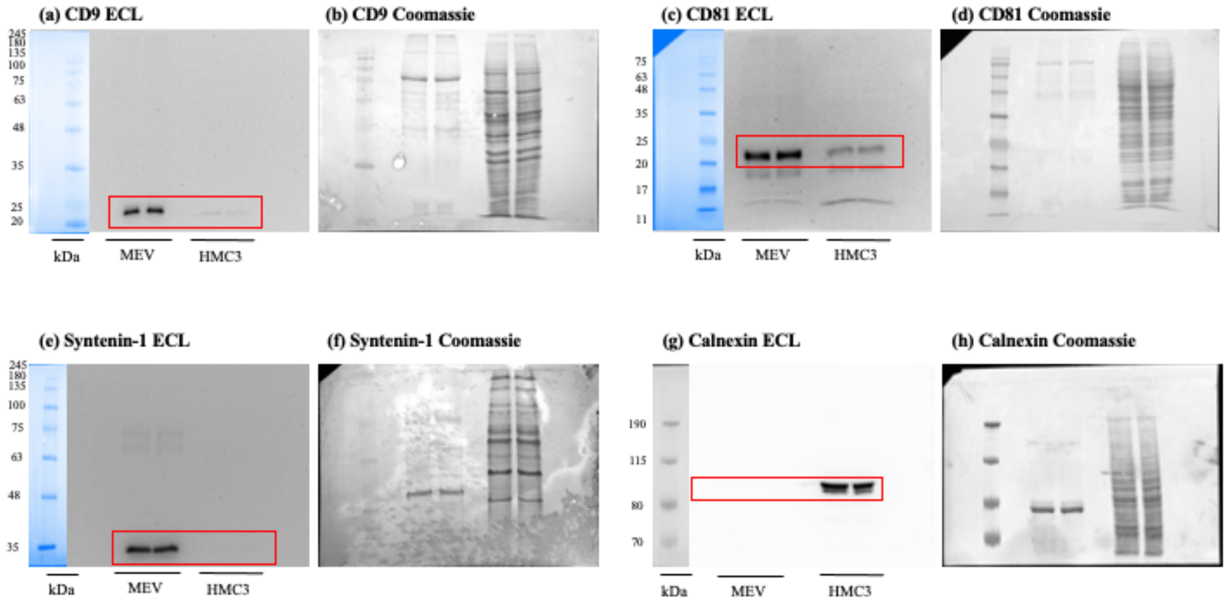


**Figure S1.** ECL and Coomassie-stained immunoblot images of MEV protein biomarkers. **(a)** ECL image for CD9. **(b)** Coomassie-stained image for CD9. **(c)** ECL image for CD81. **(d)** Coomassie-stained image for CD81. **(e)** ECL image for syntenin-1. **(f)** Coomassie-stained image for syntenin-1. **(g)** ECL image for calnexin. **(h)** Coomassie-stained image for calnexin.

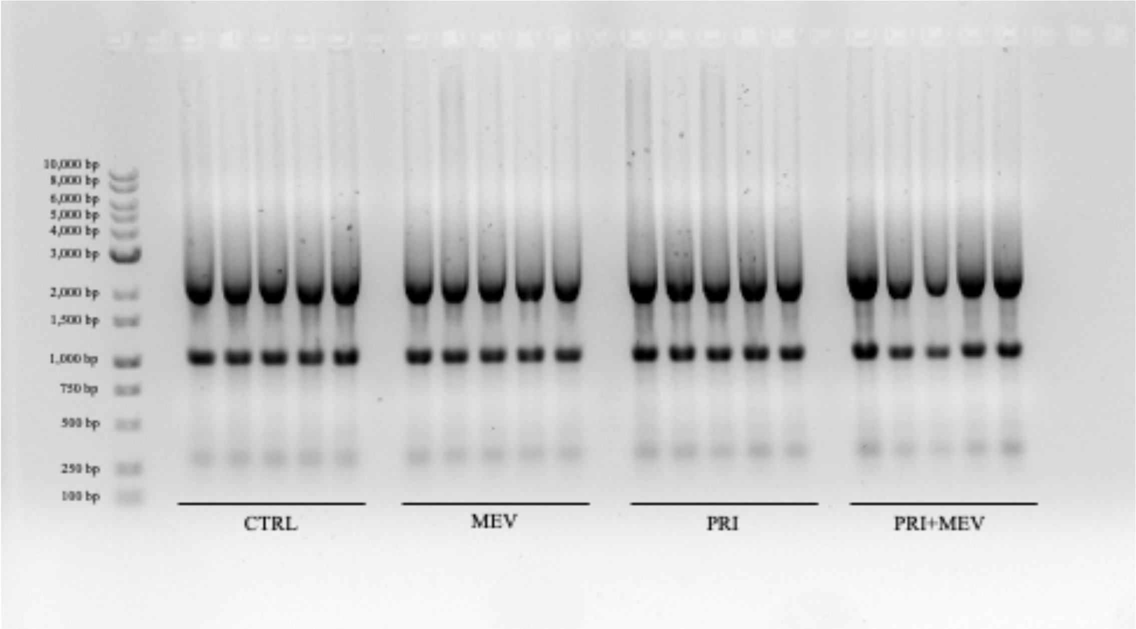


**Figure S2.** RNA profiles for 6h HMC3 samples. 1% TAE-agarose gel for verification of RNA stability and integrity. The ladder range is from 100 bp to 10,000 bp. RNA profiles for each group (CTRL, MEV, PRI, PRI+MEV) show distinct 5S, 18S, and 28S subunit distribution.

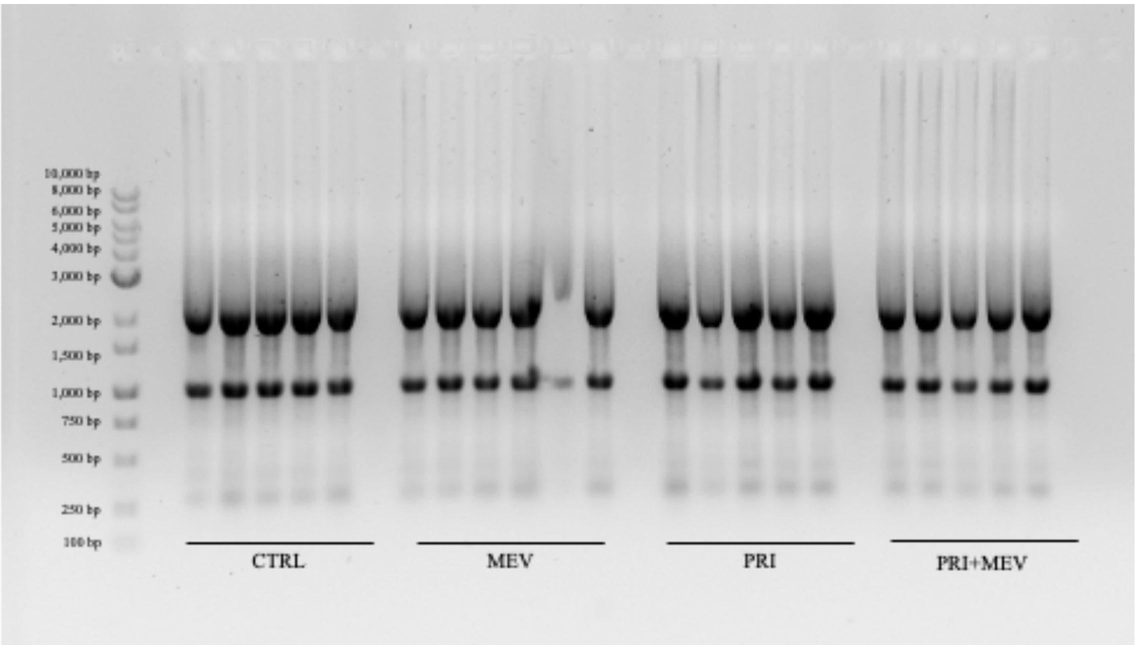


**Figure S3.** RNA profiles for 12h HMC3 samples. 1% TAE-agarose gel for verification of RNA stability and integrity. The ladder range is from 100 bp to 10,000 bp. RNA profiles for each group (CTRL, MEV, PRI, PRI+MEV) show distinct 5S, 18S, and 28S subunit distribution.

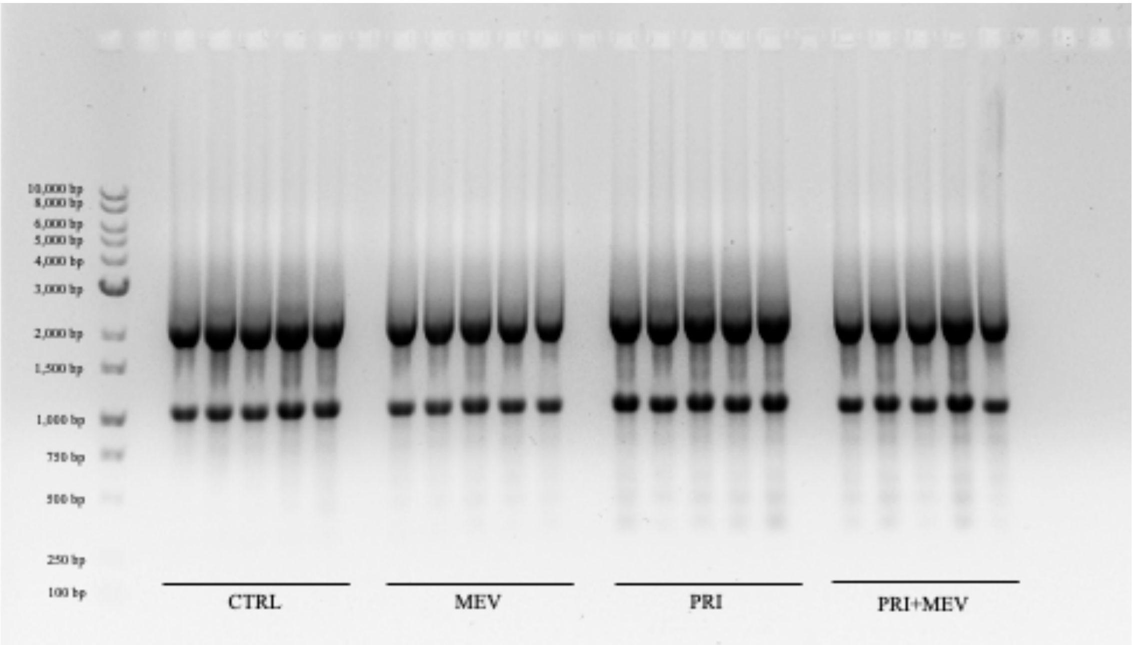


**Figure S4.** RNA profiles for 24h HMC3 samples. 1% TAE-agarose gel for verification of RNA stability and integrity. The ladder range is from 100 bp to 10,000 bp. RNA profiles for each group (CTRL, MEV, PRI, PRI+MEV) show distinct 5S, 18S, and 28S subunit distribution.

**Table S3. Primer sequences and RT-qPCR parameters for HSR genes in the HMC3 model**. Targets are identified by gene name, NCBI accession number, forward and reverse primer sequences, annealing temperature (°C), cDNA amount (ng), and their protein product. The sources of the primers (designed using NCBI or obtained from literature) are indicated. All primers target *Homo sapiens*.

| Gene | NCBI Accession # | Forward primer (5'–3') | Reverse primer (5'–3') | Annealing temperature (°C) | cDNA loaded (ng) | Primer source |
|---|---|---|---|---|---|---|
| *18S rRNA* | XR_007090847.1 | ATGGTAGTCGCCGTGCCTA | CTGCTGCCTTCCTTGGATG | 60 | 10 | Designed |
| *GAPDH* | NM_002046.7 | ATCACTGCCACCCAGAAGAC | ACCTGGTGCTCAGTGTAGCC | 63 | 10 | (Li et al. 2009) |
| *PKM* | XM_054378188.1 | GGTTCGGAGGTTTGATGA | GGCTTCTTGATCATGCTCT | 63 | 10 | (Hernández-Ochoa et al. 2021) |
| *HSF1* | NM_005526.4 | CATGAGCCTGCCTGACCTTG | CACCAGCTGCTTCCCTGAATC | 58 | 10 | (Padhy et al. 2017) |
| *HSPB1* | NM_001540.5 | CACGCAGTCCAACGAGATCA | GCGGCAGTCTCATCGGATTT | 60 | 10 | Designed |
| *DNAJB1* | NM_001539.4 | GCTGCAACGGAAGGAAGATA | TGATAATATCGCCTGGCTCC | 60 | 10 | (Chand et al. 2021) |
| *HSPA1A* | NM_005345.6 | ACATCAGCCAGAACAAGCGA | AGTCGATGCCCTCAAACAGG | 60 | 20 | Designed |
| *HSP90AA1* | NM_005348.4 | TATTTGTCCCACGACGTGCT | ATCCTCCGAGTCTACCACCC | 60 | 10 | (Hammad et al. 2022) |

**Table S4. Primer parameters for RT-qPCR in the HMC3 model**. Targets are identified by gene name, NCBI accession number, and forward and reverse primer sequences. Product size (bp), forward and reverse melting temperatures (°C) and GC content (%), hairpin temperature (°C), and homodimer and heterodimer values (%) are listed. The sources of the primers (designed using NCBI or obtained from literature) are indicated. All primers target *Homo sapiens*.

| Gene | Product size (bp) | Melting temperature (°C) | GC content (%) | Hairpin temperature (°C) | Homodimer (%) | Heterodimer (%) |
|---|---|---|---|---|---|---|
| *18S rRNA* | 107 | 65.00 | 57.90 | 28.0 | 9.30 | 8.10 |
| *GAPDH* | 302 | 63.90 | 55.00 | 53.9 | 9.60 | 18.20 |
| *PKM* | 191 | 59.00 | 50.00 | 18.0 | 10.40 | 14.40 |
| *HSF1* | 121 | 61.03 | 60.00 | 18.5 | 13.60 | 11.52 |
| *HSPB1* | 93 | 60.39 | 55.00 | 43.0 | 12.10 | 16.38 |
| *DNAJB1* | 138 | 57.98 | 50.00 | 21.6 | 18.04 | 11.91 |
| *HSPA1A* | 123 | 59.96 | 50.00 | 21.5 | 9.42 | 13.18 |
| *HSP90AA1* | 158 | 59.68 | 50.00 | 35.3 | 17.81 | 9.30 |

**Table S5. Western immunoblotting parameters for HSR protein targets in the HMC3 model**. Targets are identified with the corresponding molecular weight (kDa), SDS-PAGE gel percentage (%), amount of protein loaded (µg), SDS-PAGE gel run time and voltage, transfer settings (time, voltage, amperage), casein blocking (percentage, time), primary antibody incubation parameters (concentration, time, temperature), and secondary antibody incubation parameters (concentration, time, temperature).

| Target | Molecular weight (kDa) | Gel % | Protein loaded (µg) | SDS-PAGE run | Transfer settings | Casein blocking | Primary antibody incubation | Secondary antibody incubation |
|---|---|---|---|---|---|---|---|---|
| HSF1 | 82 | 8 | 30 | 80 min, 180 V | 40 min, 25 V, 2.5 A | 1%, 15 min | 1:1000, 3h, 22 °C | 1:10,000, 45 min, 22 °C |
| Hsp40 | 40 | 10 | 15 | 80 min, 180 V | 30 min, 25 V, 2.5 A | 5%, 60 min | 1:1000, 24h, 4 °C | 1:10,000, 45 min, 22 °C |
| Hsp70 | 72, 73 | 8 | 15 | 70 min, 180 V | 15 min, 25 V, 2.5 A | 1%, 30 min | 1:1000, 24h, 4 °C | 1:10,000, 45 min, 22 °C |
| Hsp90 | 90 | 6 | 5 | 65 min, 180 V | 15 min, 25 V, 2.5 A | 5%, 30 min | 1:1000, 24h, 4 °C | 1:10,000, 45 min, 22 °C |
| Hsp27 | 25 | 15 | 15 | 80 min, 180 V | 40 min, 25 V, 2.5 A | 5%, 30 min | 1:1000, 24h, 4 °C | 1:10,000, 45 min, 22 °C |
| Histone H3 | 15 | 15 | 10 | 70 min, 180 V | 10 min, 25 V, 1.5 A | 1%, 30 min | 1:1000, 24h, 4 °C | 1:10,000, 45 min, 22 °C |

**Table S6. Primary antibodies used for HSR protein targets in the HMC3 model**. Information includes the commercial antibody name, protein target, purchasing company and catalogue number, antigen species targeted, molecular weight, clonality, and host isotype. All antibodies are cross-reactive and recognize *Homo sapiens* and *Rattus norvegicus*.

| Antibody name | Target | Company | Catalogue number | Antigen species | Molecular weight (kDa) | Clonality | Host Isotype |
|---|---|---|---|---|---|---|---|
| HSF1 Antibody | HSF1 | Cell Signalling | #4356 | Human | 82 | Polyclonal | Rabbit IgG |
| HSP27 Recombinant Rabbit Monoclonal Antibody (JJ09-13) | Hsp27 | Invitrogen | #MA5-32473 | Human | 25 | Monoclonal | Rabbit IgG |
| HSP40 (C64B4) Rabbit mAb | Hsp40 | Cell Signalling | #4871 | Human | 40 | Monoclonal | Rabbit IgG |
| HSP70 Antibody | Hsp70 | Cell Signalling | #4872 | Human | 72, 73 | Polyclonal | Rabbit IgG |
| HSP90 (C45G5) Rabbit mAb | Hsp90 | Cell Signalling | #4877 | Human | 90 | Monoclonal | Rabbit IgG |
| Histone H3 Polyclonal Antibody | Histone H3 | Invitrogen | #PA5-31954 | Human | 15 | Polyclonal | Rabbit IgG |

**(i) HSF1 6h ECL and Coomassie images**

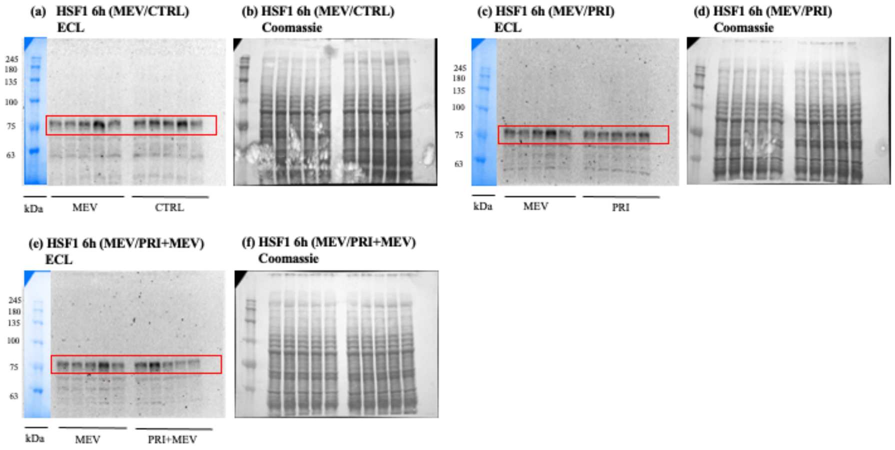


**(ii) HSF1 12h ECL and Coomassie images**

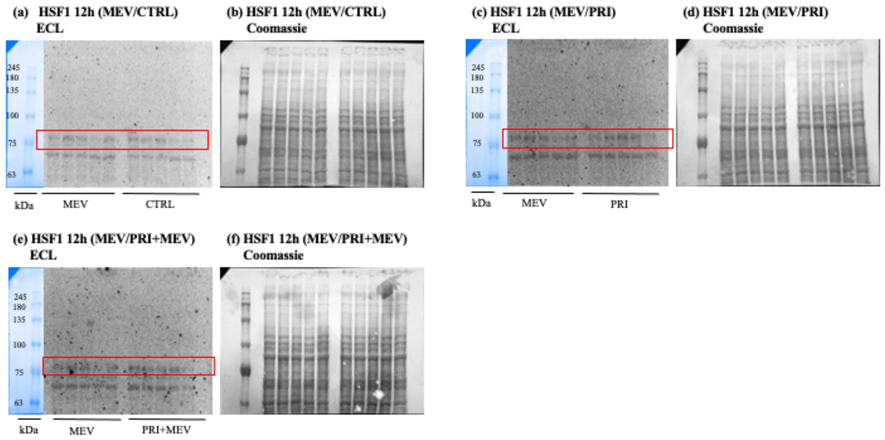


**(iii) HSF1 24h ECL and Coomassie images**

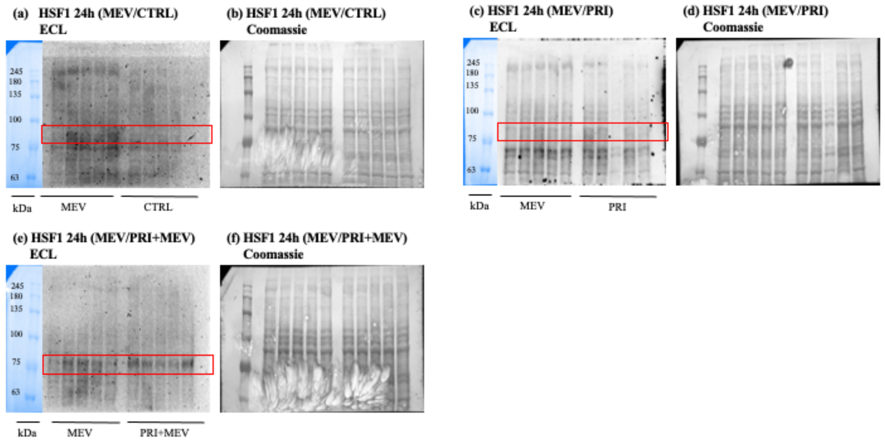


**Figure S5.** ECL and Coomassie-stained immunoblot images of HSF1 in HMC3. **(i)** ECL and Coomassie-stained images for HSF1 at 6h. **(ii)** ECL and Coomassie-stained images for HSF1 at 12h. **(iii)** ECL and Coomassie-stained images for HSF1 at 24h.

### (i) Hsp70 6h ECL and Coomassie images

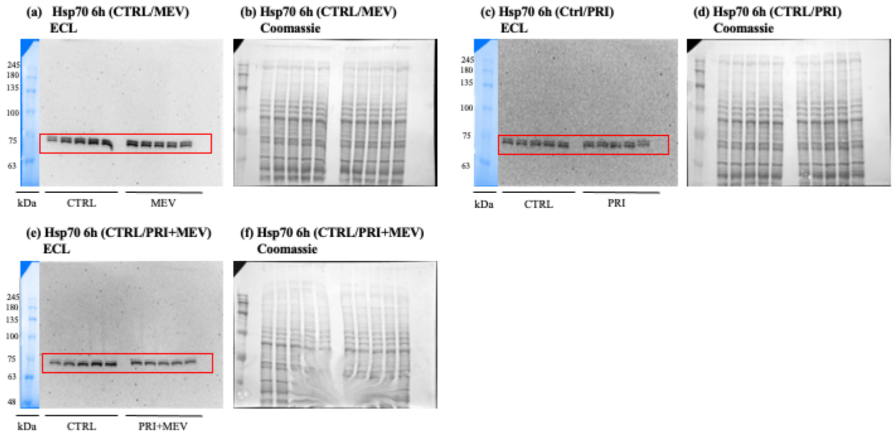


### (ii) Hsp70 12h ECL and Coomassie images

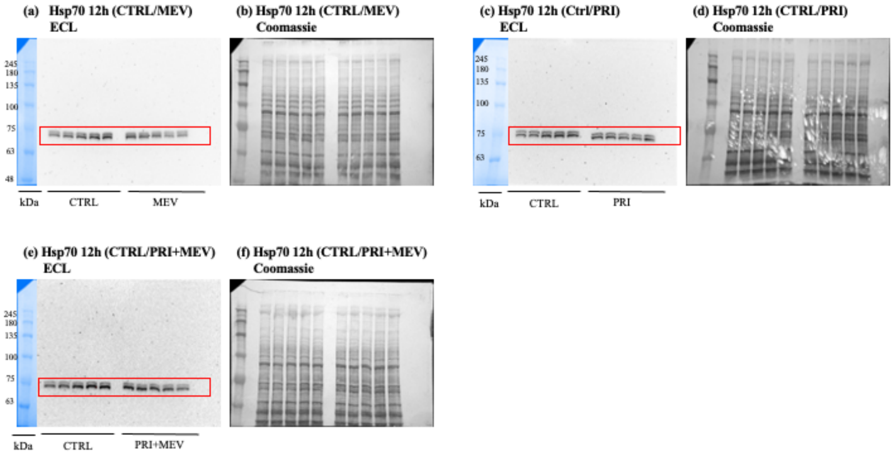


### (iii) Hsp70 24h ECL and Coomassie images

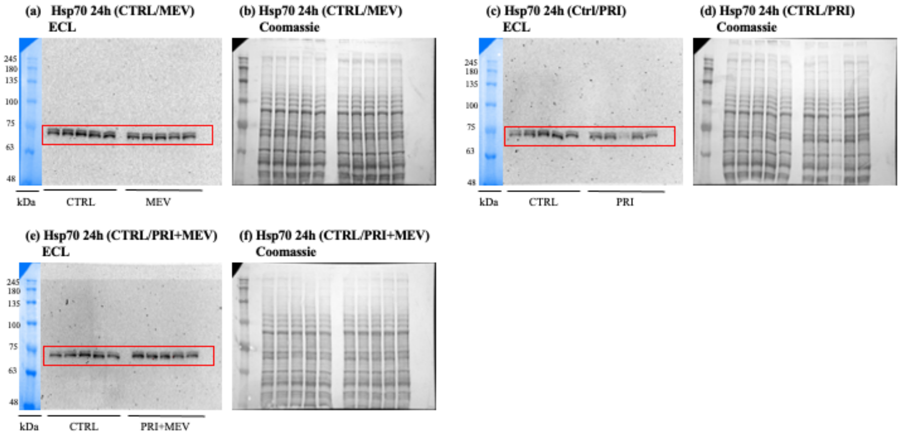


**Figure S6.** ECL and Coomassie-stained immunoblot images of Hsp70 in HMC3. (i) ECL and Coomassie-stained images for Hsp70 at 6h. (ii) ECL and Coomassie-stained images for Hsp70 at 12h. (iii) ECL and Coomassie-stained images for Hsp70 at 24h.

### (i) Hsp90 6h ECL and Coomassie images

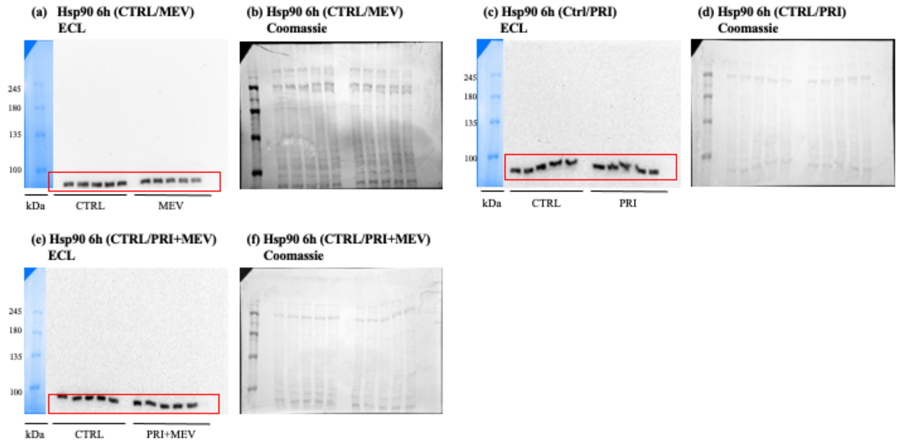


### (ii) Hsp90 12h ECL and Coomassie images

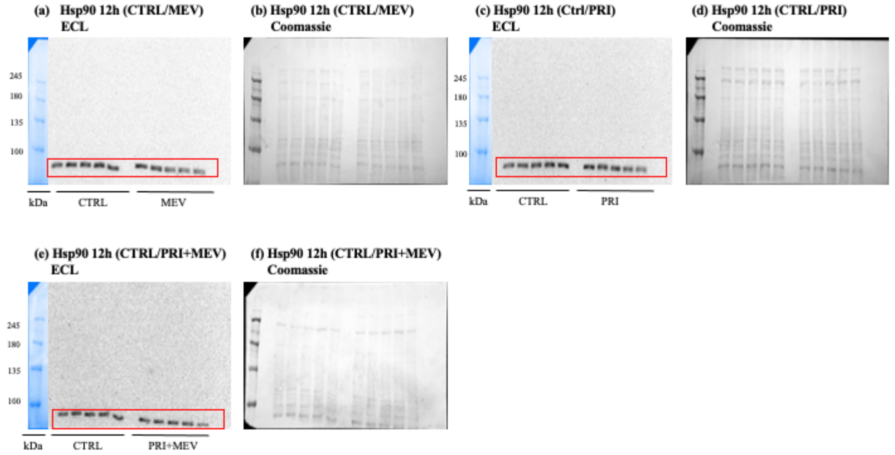


### (iii) Hsp90 24h ECL and Coomassie images

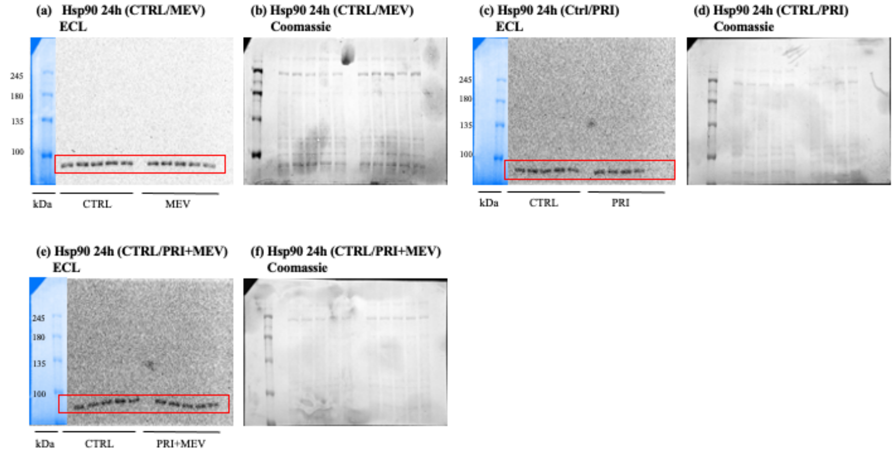


**Figure S7.** ECL and Coomassie-stained immunoblot images of Hsp90 in HMC3. (i) ECL and Coomassie-stained images for Hsp90 at 6h. (ii) ECL and Coomassie-stained images for Hsp90 at 12h. (iii) ECL and Coomassie-stained images for Hsp90 at 24h.

**(i) Hsp40 6h ECL and Coomassie images**

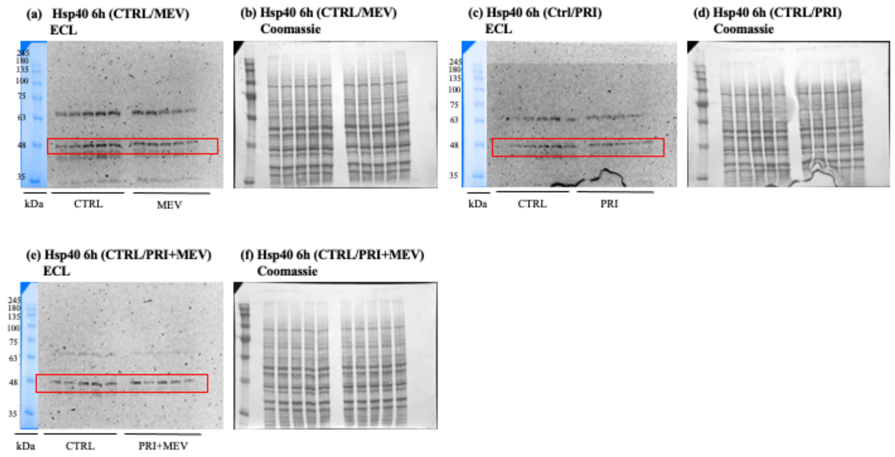


**(ii) Hsp40 12h ECL and Coomassie images**

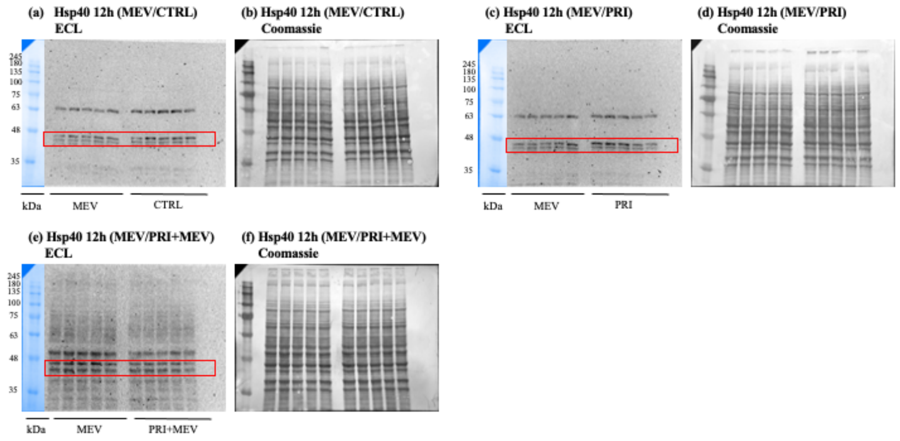


**(iii) Hsp40 24h ECL and Coomassie images**

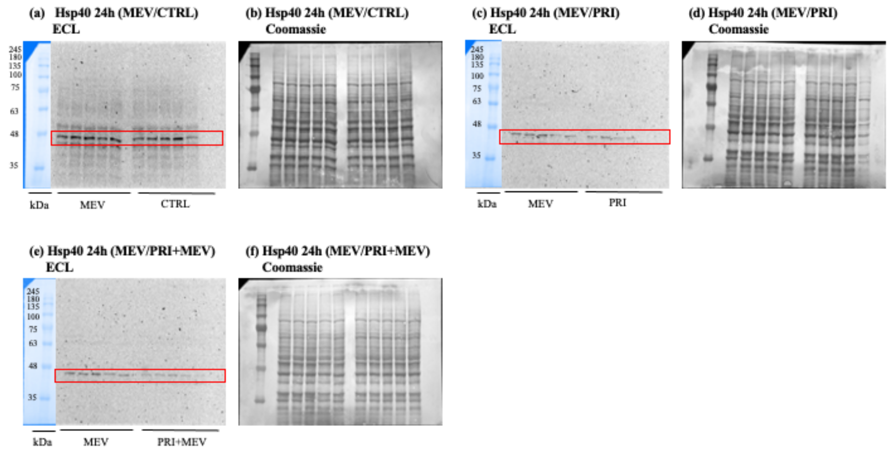


**Figure S8.** ECL and Coomassie-stained immunoblot images of Hsp40 in HMC3. (i) ECL and Coomassie-stained images for Hsp40 at 6h. (ii) ECL and Coomassie-stained images for Hsp40 at 12h. (iii) ECL and Coomassie-stained images for Hsp40 at 24h.

**(i) Hsp27 6h ECL and Coomassie images**

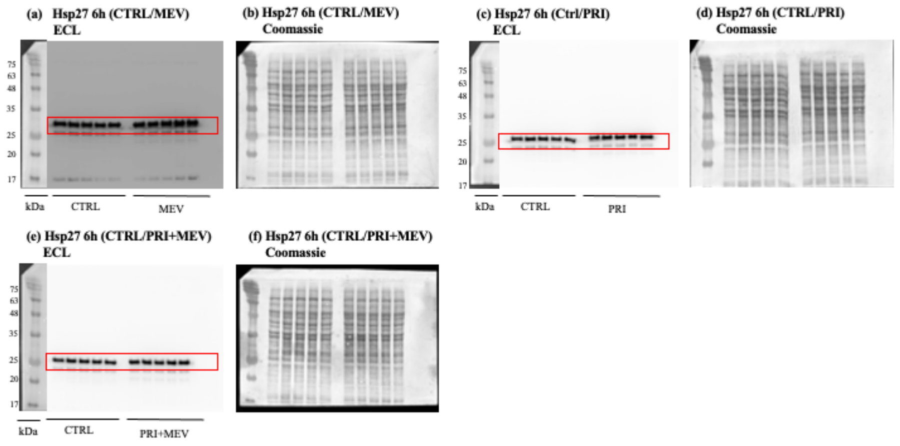


**(ii) Hsp27 12h ECL and Coomassie images**

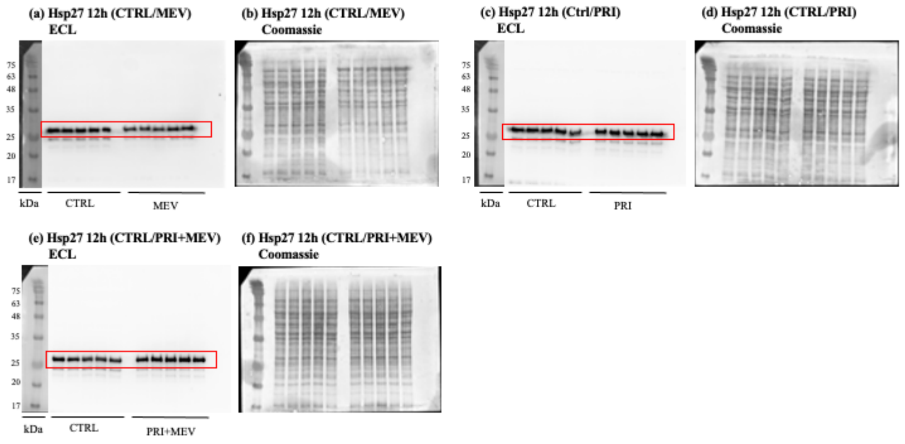


**(iii) Hsp27 24h ECL and Coomassie images**

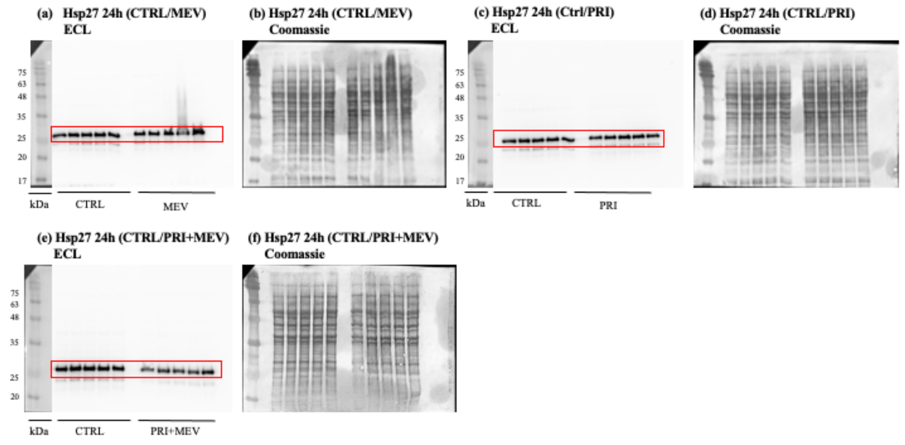


**Figure S9.** ECL and Coomassie-stained immunoblot images of Hsp27 in HMC3. (i) ECL and Coomassie-stained images for Hsp27 at 6h. (ii) ECL and Coomassie-stained images for Hsp27 at 12h. (iii) ECL and Coomassie-stained images for Hsp27 at 24h.
